# Mapping of multiple neurotransmitter receptor subtypes and distinct protein complexes to the connectome

**DOI:** 10.1101/2023.10.02.560011

**Authors:** Piero Sanfilippo, Alexander J. Kim, Anuradha Bhukel, Juyoun Yoo, Pegah S. Mirshahidi, Vijaya Pandey, Harry Bevir, Ashley Yuen, Parmis S. Mirshahidi, Peiyi Guo, Hong-Sheng Li, James A. Wohlschlegel, Yoshinori Aso, S. Lawrence Zipursky

**Author notes:** Correspondence (S.L.Z.).

## Abstract

Neurons express different combinations of neurotransmitter receptor (NR) subunits and receive inputs from multiple neuron types expressing different neurotransmitters. Localizing NR subunits to specific synaptic inputs has been challenging. Here we use epitope tagged endogenous NR subunits, expansion light-sheet microscopy, and EM connectomics to molecularly characterize synapses in Drosophila. We show that in directionally selective motion sensitive neurons, different multiple NRs elaborated a highly stereotyped molecular topography with NR localized to specific domains receiving cell-type specific inputs. Developmental studies suggested that NRs or complexes of them with other membrane proteins determines patterns of synaptic inputs. In support of this model, we identify a transmembrane protein associated selectively with a subset of spatially restricted synapses and demonstrate through genetic analysis its requirement for synapse formation. We propose that mechanisms which regulate the precise spatial distribution of NRs provide a molecular cartography specifying the patterns of synaptic connections onto dendrites.

## Introduction

Recent progress in electron microscopy (EM) has enabled generation of single-synapse level connectomes of large brain volumes (MICrONS Consortium, 2023; Scheffer et al., 2020; Shapson-Coe et al., 2021; Shinomiya et al., 2019; Takemura et al., 2013; Winding et al., 2023). In *Drosophila*, EM reconstructions revealed extraordinary complexity and specificity of brain wiring. Hundreds of neuron types form specific patterns of connections with multiple partners in highly reproducible ways (Scheffer et al., 2020; Takemura et al., 2015). Single-cell sequencing has uncovered distinct patterns of mRNAs in different neurons for proteins involved in the development and function of synapses (Kurmangaliyev et al., 2020; Özel et al., 2021). This is particularly striking in the transcriptional expression patterns of neurotransmitter receptor (NR) subunits.

NRs fall into three families: pentameric cys-loop ionotropic receptors, tetrameric glutamate ionotropic receptors, and G protein-coupled metabotropic receptors. In this study, we focus on NRs of the cys-loop superfamily. Cys-loop receptors mediate responses to the major excitatory and inhibitory neurotransmitters in the *Drosophila* brain, acetylcholine, and GABA, respectively.

These subunits come together in different combinations to form ligand-gated ion channels. The expression of distinct combinations of NR subunits gives rise to multiple molecularly distinct receptor complexes (Chamaon et al., 2002; Gisselmann et al., 2004; Lansdell et al., 2012; Schulz et al., 1998; Zhang et al., 1995). Different mammalian GABA receptors of this family have been shown to localize to different domains of pyramidal cells (e.g., axon initial segment vs soma) (Contreras et al., 2019; Panzanelli et al., 2011). Previous studies in *Drosophila* have also shown that targeted expression of cDNAs encoding tagged versions of the GABA receptor subunit Rdl and the nicotinic subunit nAChRα7 resulted in localization to different dendritic domains in motor neurons (Ryglewski et al., 2017) and in T4/T5 neurons in the visual system (Fendl et al., 2020). Voltage and calcium imaging in visual circuits in the fly revealed characteristic patterns of activity in different neuronal compartments (Yang et al., 2016). It remains largely unknown what underlies these subcellular specific activity patterns at the molecular level. They likely include, in part, the subunit composition of NRs in the postsynaptic membrane and their spatial distribution in dendrites.

Localizing NRs at synapses in dendrites has been problematic for two reasons. First, it has been difficult to generate antibodies for multi-pass transmembrane proteins (Hamakubo et al., 2014), which hinders specific immunohistochemical labeling of many NR subunits. To address this issue, we used CRISPR-modification to introduce various epitope tags into the endogenous NR loci. These modified loci allowed us to label NR subunits with highly specific commercially available antibodies while preserving their endogenous expression patterns. We also engineered conditional alleles for the selective labeling of tagged NR subunits in single neurons through cell-type-specific expression of recombinases. Second, the density of the neuropil and diffraction-limited light microscopy preclude localizing receptors to synapses in identified neurons. And third, there are substantial technical challenges of EM localization of proteins (Boassa, 2015). To overcome these limitations, we take advantage of recent developments in Expansion Microscopy to localize protein in isotropically expanded tissue (Chen et al., 2015) using custom-built lattice light-sheet (Chen et al., 2014a; Gao et al., 2019) and commercially available light-sheet microscopy (ExLLSM and ExLSM, respectively), to achieve effective super-resolution.

Here, we localize tagged NR subunits in specific neuron types throughout the brain in adult and developing neurons at the level of single cells, and at single synapses between identified neuron types at super-resolution. We focus on the distribution of seven different subunits in directionally selective motion sensing neurons and demonstrate that these NRs are localized to specific spatial domains along the proximodistal dendritic axis. Using affinity-purification-mass-spectrometry, we identify a transmembrane protein implicated in synaptic adhesion associated with an NR subunit selectively localized to one of these domains. Our findings raise the possibility that NR protein complexes selectively localized to specific dendritic domains provide molecular cues specifying patterns of synaptic inputs from different neuron types.

## Results

### NR subunits are expressed in different brain regions

There are around 60 genes encoding subunits of NRs in the *Drosophila* genome and about 100 in the mouse and human genomes (Deng et al., 2019; Kondo et al., 2020; Littleton and Ganetzky, 2000). We generated tagged alleles of 11 NR subunits of the cys-loop superfamily, which form pentameric ligand-gated ion channels. These include receptors responsive to acetylcholine, γ-aminobutyric acid (GABA) and glutamate, the three major neurotransmitters in the fly (Figure 1A; Table S1). Endogenous tagging of NRs is preferrable to transgenic expression systems for studying NR localization, as the endogenous proteins preserve their unique, cell-type specific patterns and developmental expression (Fendl et al., 2020; Mikuni et al., 2016).

**Figure 1.**
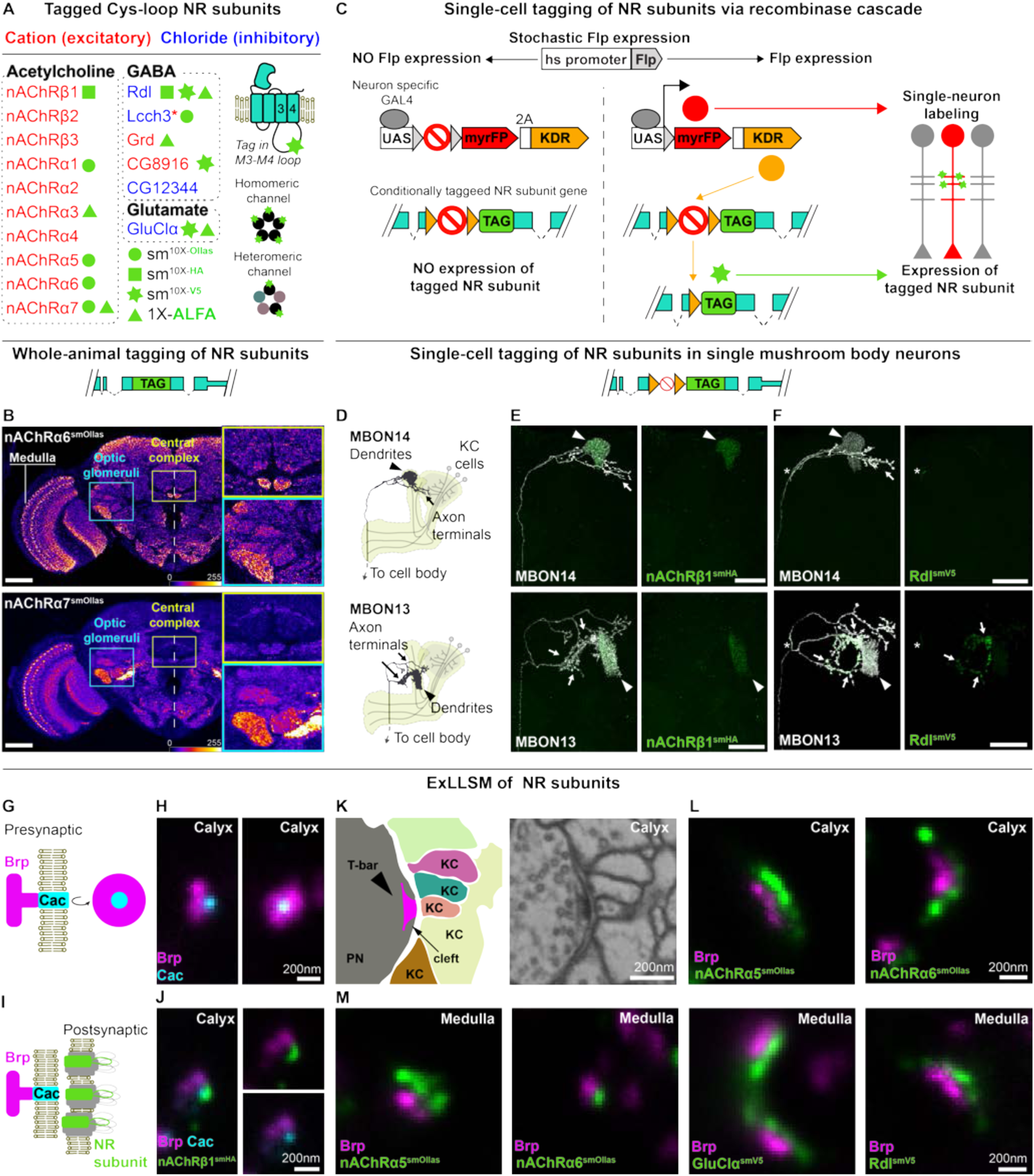
Tagged NR subunits localize to synapses. (A) Cys-loop superfamily NR subunits tagged in this study. For some NRs, two or three different tagged versions were made as indicated. Red asterisk, Lcch3 forms cationic channels with Grd and with CG8916 and chloride channels with Rdl (Gisselmann et al., 2004; Huang et al., 2021; Zhang et al., 1995). The epitope tag was inserted into a poorly conserved region of the M3-M4 cytoplasmic loop in each NR subunit (Figures S1A and S1B). These proteins form homo- or heteropentameric channels. (B) Whole animal tagging of NR subunits. For each conditionally tagged allele a corresponding whole-animal tagged version was generated in which all cells expressing the receptor express a tagged version of it. Expression pattern of endogenously tagged alleles of nAChRα6 (top panel) and nAChRα7 (bottom panel). Dotted line, brain midline; yellow inset, neuropils of the central complex; cyan inset, optic glomeruli. Neurons that form synapses in the medulla neuropil of the optic lobe as indicated are the focus of this study. Brp staining visualizes neuropils. Scale bars, 50 μm. (C) Schematic for conditional tagging of NR subunits in sparsely distributed neurons. See Figure S3B for details. (D-F) Localization of NR subunits to domains of neurons with complex morphology in the central brain. MBONs have two main processes, one which terminates in a compact dendrite and a second axonal process, which makes secondary projections to distinct brain areas. (D) Morphology of MBON14 or MBON13 – dendrites innervate select compartments of the mushroom body (yellow). Kenyon cells provide cholinergic inputs to MBON dendrites. Localization of nAChRβ1-smHA (E) or Rdl-smV5 (F) in MBON14 or MBON13. Arrowhead – dendrites; arrows – axonal projections; asterisk – putative axon initial segment. Scale bars, 25 μm. (G) Schematic of synaptic active zone marked by Brp and the voltage-gated calcium channel Cacophony (Cac). In EM, cytoplasmic Brp protein localizes to a presynaptic T-bar structure associated with the presynaptic membrane (see panel K) and as a donut by STED and LLSM when immunostained for the Brp-directed antibody nc82. (H) Brp with Cac at its center visualized using ExLLSM. The synapse shown is from the mushroom body calyx in the central brain. Lateral (left) and planar (right) views are shown. (I) Schematic of NR subunits juxtaposing Brp and Cac. (J) nAChRβ1 subunits cluster juxtaposed to an active zone in the calyx. Lateral views are shown. (K) EM (right) and schematic (left) of multiple-contact synapse between projection neurons (PN) and Kenyon cells (KC) in the mushroom body (single synapse from EM data from Zheng et al., 2018). (L) Immunostaining of PN-KC synapses, as indicated. (M) Examples of active zones and different NR subunits (as indicated) paired at synapses in the medulla neuropil. Scale bars in H, J, L and M panels represent unexpanded tissue size (adjusted for 4.65X expansion factor, see Methods).

To generate tagged alleles, the endogenous genomic loci encoding NR subunits were modified by directed knock-in of DNA sequences encoding epitope tags using CRISPR-targeted recombination (Gratz et al., 2015; Kanca et al., 2019) (Figure S1A). All tags were inserted into the cytoplasmic loop between the M3 and M4 transmembrane domains, which are shared amongst all cys-loop receptors (Figures 1A and S1B). Several studies have shown that NRs with insertions in this loop are functional and localize to postsynaptic sites (Drenan et al., 2008; Nashmi et al., 2003; Raghu et al., 2009; Slimko and Lester, 2003). NR subunits form homomeric or heteromeric NRs, or both (Figure 1A), resulting in a large array of distinct NR types with unique functional properties (Boorman et al., 2003; Gisselmann et al., 2004; Lansdell et al., 2012).

All tagged NRs localized to the neuropil (Figures 1B and S1C). Their expression patterns were similar to those seen with antibodies to the unmodified NR subunits (Figure S1D). Epitope-tagged proteins rescued lethality in cases where mutant alleles were not viable (see Methods). Furthermore, all remaining homozygous tagged alleles were viable. Typically, the tagged NR subunits were broadly expressed throughout the brain, with some subunits highly enriched in specific neuropils. Each NR subunit exhibited a characteristic pattern of expression (Figures 1B and S1C-S1D; also see Figure S3A).

Due to the density of processes within neuropils, it is not possible to assign expression of proteins detected by immunofluorescence to specific neuron types. To overcome this limitation, we generated inducible alleles and a new approach to tag NR subunits in identified cell types with single cell resolution (Figures 1C, S1A and S3B). Reagents to label virtually any neuron in *Drosophila* in this way are available (Jenett et al., 2012; Pfeiffer et al., 2010). In this study, we explored the distribution of NR subunits in different neurons in the visual system and the mushroom body.

The mushroom body is the center for associative learning in insect brains and its sensory inputs and output synapses are cholinergic (Figure S2A) (Barnstedt et al., 2016; Yasuyama et al., 2002). As expected from RNA-seq data (Aso et al., 2019), we observed the nAChRβ1 subunit in the dendrites of two different MBONs (Figures 1D and 1E). In contrast, the subcellular distribution of the GABAergic subunit Rdl in MBONs was highly cell-type-specific (Figures 1D, 1F and S2C-S2D). For instance, in MBON05 Rdl localized to both axons and dendrites while it was predominantly restricted to dendrites in MBON11 (Figures S2C and S2D). Thus, tagged NR subunits localize to specific neuronal domains and these may differ even between neurons with closely related functions. We next sought to address whether the tagged NR subunits localized to synapses.

### Tagged NR subunits localize to synapses

The resolution of light microscopy is not sufficient to localize NRs to synapses. To increase the effective resolution, we imaged expanded tissue (Chen et al., 2015) using a lattice light-sheet microscope (ExLLSM) (Chen et al., 2014a). This method allows imaging of large volumes at an effective resolution of approximately 60 by 60 by 90 nm (Gao et al., 2019). We assessed the localization of the tagged NRs by comparing images of synapses in the mushroom body with those previously obtained using stimulated emission depletion microscopy (STED) and EM.

Previous STED studies of the mushroom body (MB) calyx identified presynaptic sites characterized by Brp-stained donut-shaped structures with the Ca2+ channel Cacophony (Cac) at the center (Fulterer et al., 2018) (Figure 1G). Brains bearing an ALFA-tagged allele of Cac and stained for both the ALFA tag and Brp were imaged. We observed presynaptic structures in the MB calyx and in the medulla region of the optic lobe of similar dimensions to those previously described in the MB calyx in STED and EM studies (Figures 1G-1J and S1E). Tagged nAChRβ1 clustered in juxtaposition to these presynaptic sites (Figures 1I-1J and S1F). The synaptic structures identified by ExLLSM for other NR subunits in the calyx and medulla were also similar (Figures 1K-1M and S1G-S1I). All NR subunits were preferentially associated with Brp (see Figures S2M-S2P for quantification), consistent with their synaptic localization. In addition, tagged nAChR subunits localized in different types of synaptic structures similar to those described by EM (Figures 1K, S2A-S2B and S2E-S2L) (Fulterer et al., 2018; Takemura et al., 2017a; Yasuyama et al., 2002).

In summary, endogenously tagged NRs visualized by ExLLSM were distributed in NR-type-specific patterns within the CNS and selectively localized to postsynaptic sites consistent with previous studies using EM and STED.

### NR subunit distribution matches the EM connectome

We next sought to determine whether the patterns of NRs correlated with the distribution of inputs from cholinergic, GABAergic, and glutamatergic neurons as determined by connectomic and gene expression studies. We focused on circuits in the fly visual system and, in particular, the medulla neuropil. The repetitive pattern allowed us to look at many different neurons of the same type in a single animal, and it is straight forward to generate sparsely labeled neurons of the same type. In the medulla neuropil overlapping processes of many different neuron types form stereotyped circuits (Figure 2A). Each neuron type expresses different levels and combinations of transcripts encoding NR subunits (Figure S3A) (Kurmangaliyev et al., 2020).

**Figure 2.**
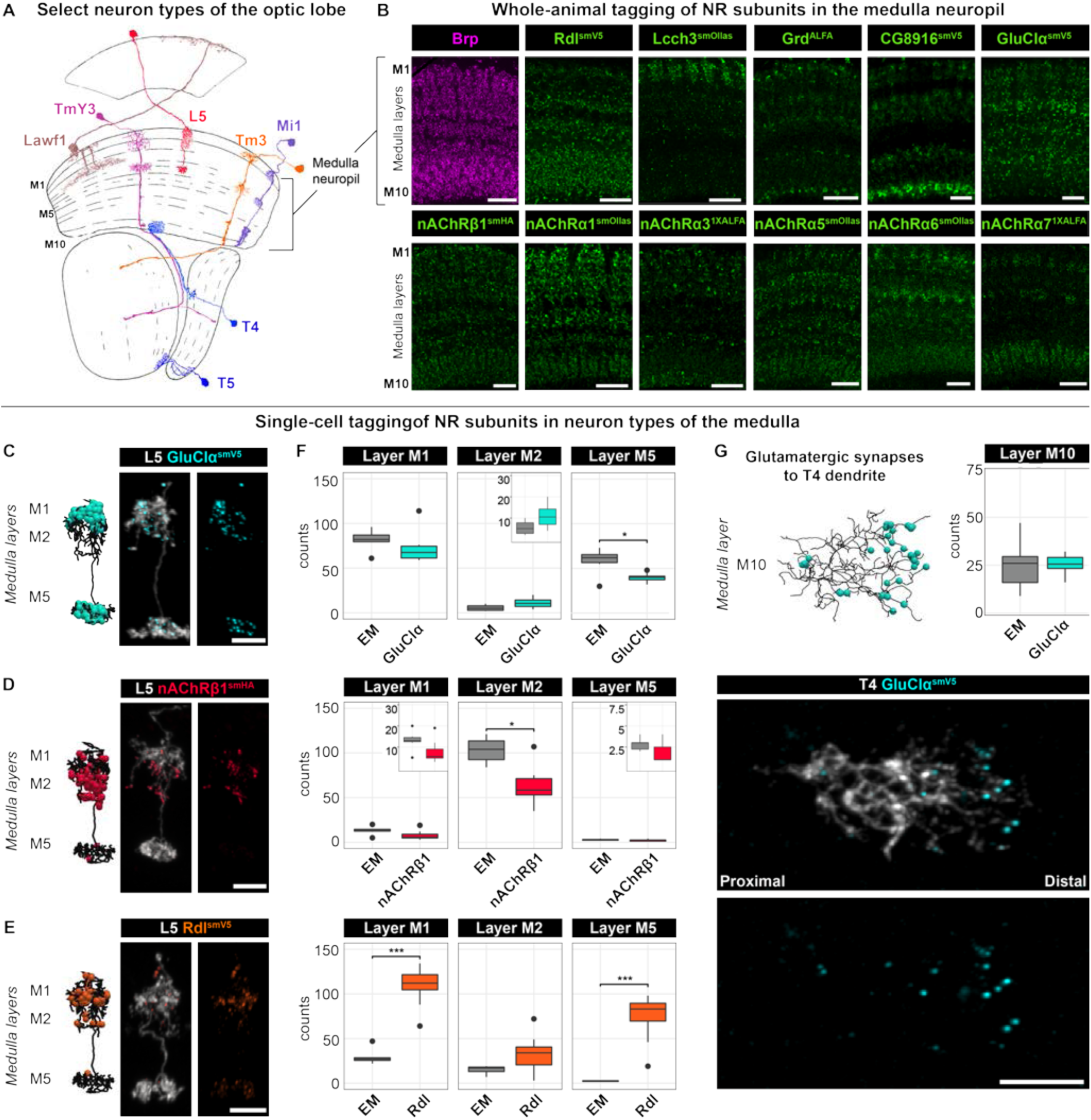
NR subunit distribution matches the EM connectome. (A) Morphologies of a small selection of neuron types of the ∼150 different types in the fly optic lobe. Adapted from (Fischbach and Dittrich, 1989). Three of the ten medulla neuropil layers are labeled (M1, M5 and M10). (B) NR subunit expression in medulla layers. The presynaptic marker Brp (magenta) detected with anti-Brp antibody is seen in all 10 medulla layers (M1-M10). Distribution of NR subunits (green) in the medulla as indicated. NR subunits visualized using antibodies to the epitope tag (indicated by the superscript: anti-V5, anti-Ollas, anti-HA, or anti-ALFA). Scale bar, 10 μm. (C-E) Localization of GluClα-smV5 (C), nAChRβ1-smHA (D), and Rdl-smV5 (E) in dendrites of L5 neurons. Left, morphology of reconstruction of the EM-based inputs to L5 dendrites in the medulla and annotated synapses for each neurotransmitter type (Takemura et al., 2015). Right two panels, neuron morphology (gray) and NR subunits in color as indicated. Scale bars, 5 μm. (F) Quantification of data shown in E-G and from the EM data (EM, n=7; Rdl, n=15; GluClα, n=12; and nAChRβ1, n=8). Bonferroni adjusted p-value (*), < 0.05, (***), < 0.001 from Wilcoxon rank-sum test. See text for discussion of discrepancies between EM and tagged puncta. (G) Distribution of GluClα-smV5 in T4 dendrites. Quantification data same as in Figure S4F. Scale bars, 5 μm.

The 10 layers of the medulla comprise the highly branched processes and synapses of many neurons (>10,000 neurons, >100 neuron types and ∼2 million synapses) (Takemura et al., 2015). Some receptors are broadly expressed, whereas others are preferentially enriched in specific layers (Figure 2B). For instance, the inhibitory GABA receptor subunit Rdl, which can form homomeric and heteromeric receptors (Gisselmann et al., 2004; Zhang et al., 1995), is expressed in most layers of the medulla. By contrast, the expression of Lcch3, another GABA receptor subunit which can form heteromeric receptors with Rdl (Zhang et al., 1995) is more restricted (Figure 2B). Differential localization was also observed for different nAChRs subunits (Figure 2B).

EM-level connectomes have established that many neuron types in the medulla receive inputs from multiple types of presynaptic neurons (Takemura et al., 2015, 2017b). For instance, GluClα localized to domains of L5 which predominantly receive glutamatergic inputs in medulla layers M1 and M5 (Figure 2C). The number of puncta seen for GluClα in L5 neurons correlated well with EM data (Figure 2F). These neurons also receive cholinergic and GABAergic innervation to the same and different dendrites in discrete and reproducible patterns and this is reflected in the distribution of Rdl and nAChRβ1 (Figures 2D-2F). Discrepancies between synapse numbers were observed in some cases with Rdl and different acetylcholine receptors. This may represent NRs that are extrasynaptic (e.g., small Rdl puncta) and different receptors for the same neurotransmitter at different synapses in the same neuron (e.g., multiple nAChRs). In general, however, there was a good correlation between the distribution of receptor puncta and synapses determined by EM across several neuron types (Figures 2G, S3C-S3F and S4F).

### NR subunits are differentially localized along dendrites

The precise distribution of synaptic inputs from different neurons along the proximodistal axis of T4 dendrites is proposed to play a crucial role in motion detection. T4 dendrites receive inputs from GABAergic, cholinergic, and glutamatergic neurons from eight identified cell types in specific domains along the proximodistal axis (Figure 3A) (Shinomiya et al., 2019; Takemura et al., 2017b). The synaptic inputs to T5 are different (Figure S4A; see below), with different neuron types also forming synapses within specific dendritic domains (Shinomiya et al., 2019).

**Figure 3.**
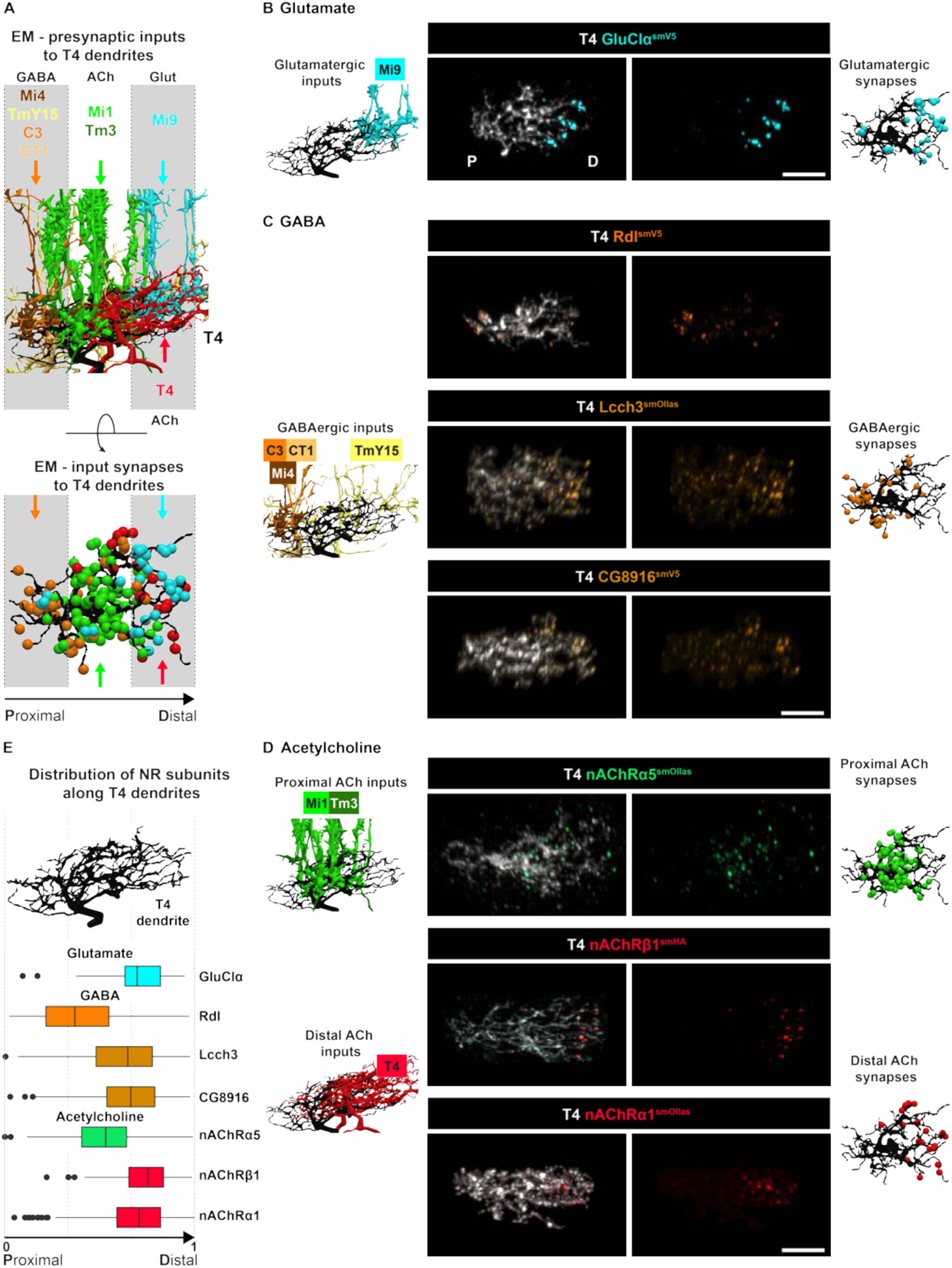
NR subunits are differentially localized along dendrites. (A) Upper panel, EM reconstruction of different presynaptic inputs, as indicated, along the proximodistal axis of T4 dendrites. Dendrites span an average of three columns. Lower panel, annotated synapses for different neurotransmitter inputs are shown. Colored arrows in the lower panel show domains targeted by color matched presynaptic inputs from the upper panel. TmY15 is an exception forming synapses across the length of T4 dendrites. (B-D) Localization of tagged NR subunits in dendrites of single T4 neurons. Left-most column, EM reconstruction of glutamatergic, GABAerbic and cholinergic inputs. Central columns, NR subunit pattern observed by conditional tagging in single T4 dendrites. Right-most column, subset of EM-annotated synapses as shown in the lower part of panel A. Scale bars, 5 μm. (E) Quantification of NR subunit expression along the normalized proximo-distal axis of T4 dendrites (See methods). Rdl, n = 6; CG8916, n=4; Lcch3, n=3; GluClα, n = 4; nAChRα1, n = 5; nAChRα5, n = 5; nAChRβ1, n = 10.

The distribution of NRs in T4 dendrites was consistent with the neurotransmitters used by the synaptic inputs. GluClα, as described above, was highly localized to the distal domains (Figures 3B and 3E). By contrast, Rdl was strongly enriched in the proximal region with additional puncta sparsely scattered throughout the arbors (Figures 3C and 3E). T4 neurons also express two additional GABA NR subunits, Lcch3 and CG8916 (see Figure S3A). These two subunits were not localized proximally, but rather exhibited a common distribution with enrichment in the distal region and then sparsely throughout the rest of the dendrite (Figures 3C and 3E). The proximal enrichment of Rdl matches the innervation pattern by three different GABAergic neuron types (i.e., Mi4, C3 and CT1). By contrast, TmY15 synapses scattered along the length of T4 dendrites may signal through NRs containing Lcch3 and CG8916.

Different nAChR subunits also show different distributions in T4 dendrites. nAChRα5 localized to the middle domain of T4 dendrites, whereas both nAChRβ1 and nAChRα1 localized to the distal domain (Figures 3D and 3E). These patterns correlate with the cholinergic inputs from overlapping Mi1 and Tm3 neurons in the middle domain, and to dendrodendritic synapses of overlapping T4 dendrites in the distal domain. These data suggest that the subunit composition of NRs to the same neurotransmitter is unique to specific synaptic inputs.

T5 dendrites receive inputs different from T4 and these also showed specificity along the proximodistal axis (Figures S4A-S4E). Rdl, nAChRα5, nAChRβ1 and nAChRα1 subunits were distributed in a fashion similar to T4 (Figures S4C-S4E). GluClα was not detected in T5 dendrites, consistent with the absence of glutamatergic inputs (Figures S4A and S4B). CG8916 and Lcch3 also lacked the enrichment to the tips observed for T4 dendrites (Figures S4C and S4E). There were twice the number of nAChRβ1 puncta in T5 as in T4 dendrites (Figure S4F). This is consistent with the additional cholinergic input T5 dendrites receive in the distal region by Tm9 neurons (Figures S4A and S4D). Three other Tm neuron types (Tm1, Tm2, and Tm4) provide inputs to the middle domain in a pattern similar to the distribution of nAChRα5 (Figures S4A and S4D).

The distribution of nAChRα1, nAChRβ1 and nAChRα5 partially overlap in the middle domain. To assess whether these nAChR subunits were in the same or different synapses, we tagged nAChRα1 or nAChRα5 in combination with nAChRβ1 in the same neuron with different epitopes (Figure S4G). nAChRα1 puncta largely co-localized with nAChRβ1, suggesting that these are found at the same synapses (Figures S4H and S4I). By contrast, nAChRα5 and nAChRβ1 did not co-localize demonstrating that these are not at the same synapse (Figures S4H and S4I). Thus, even within the same dendritic domain different cholinergic neuron types may form synapses selectively onto postsynaptic sites expressing different receptors.

In summary, the distribution of NR subunits reflected the specific arrangement of various GABAergic, cholinergic, and glutamatergic inputs onto T4 or T5 dendrites with spatial specificity along their proximodistal axis. Different neuron types, which use the same neurotransmitter, may communicate through molecularly distinct receptors within the same or different dendritic domains.

### Localization of NR subunits during development is cell-type specific

We next sought to explore how NRs become localized to synapses during development. NRs could localize directly to the discrete domains where synapses form. Alternatively, NRs could be initially uniformly distributed and subsequently stabilized at synapses, downregulated in incorrect locations or both. Analysis of whole-animal tagged NR subunits for Rdl-smV5, nAChRβ1-smHA and GluClα-smV5 showed progressive accumulation of NR subunits in the developing medulla neuropil (Figures S5A-S5D). To address if this accumulation occurs similarly in different neurons we turned to single-cell analysis in T4, T5 and L5 medulla neurons, where these NR subunits are expressed throughout pupal development (Figures 4 and S5A).

**Figure 4.**
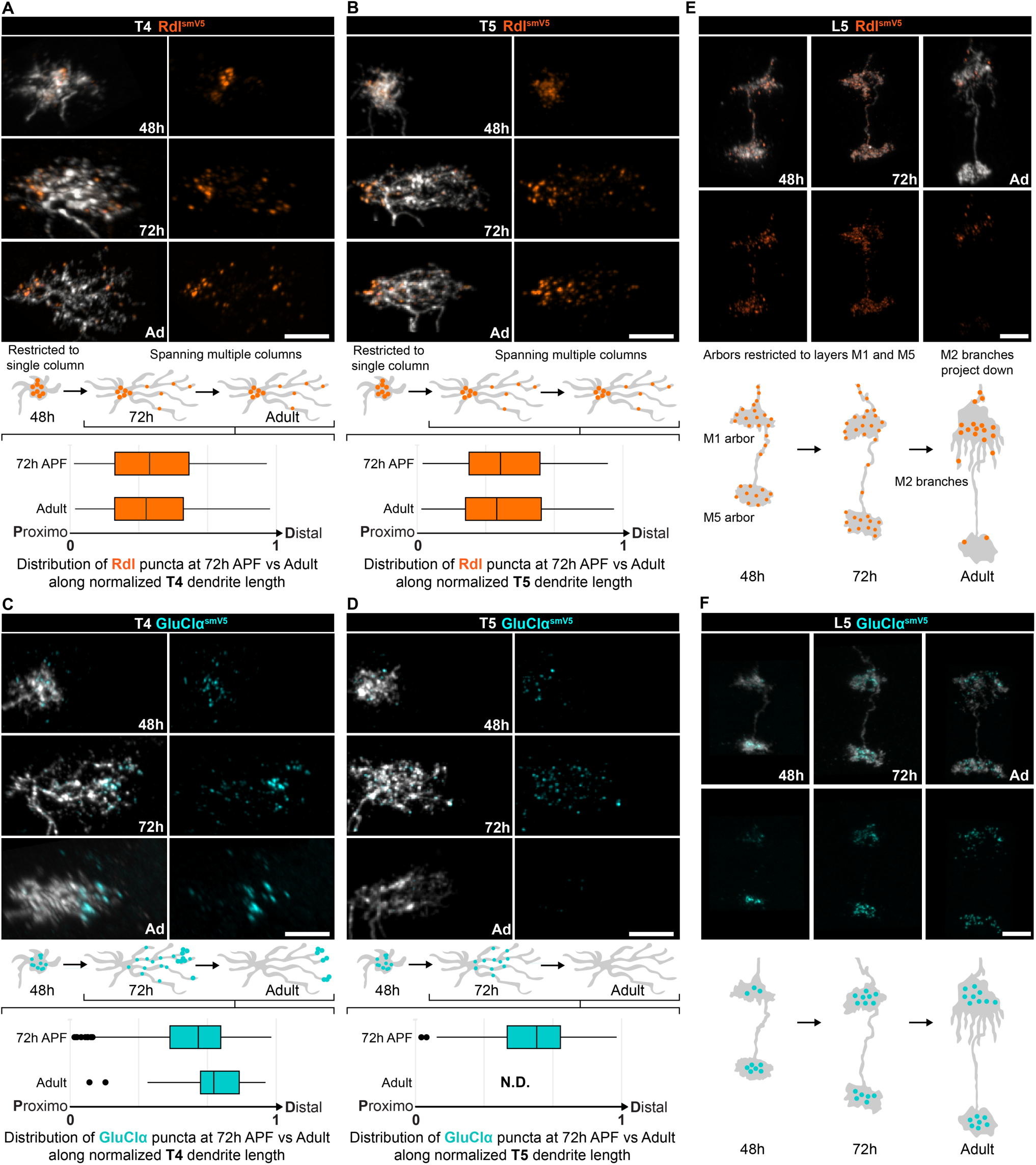
Localization of NR subunits during development is cell-type specific. Time course of NR subunit accumulation during development of T4 (A, C), T5 (B, D) and L5 dendrites (E,F) of Rdl-smV5 (A-B, E) or GluClα-smV5 (C-D, F). Times shown are 48h and 72h after pupal formation (APF) and adult (Ad). Schematic of dendrite development with NR distribution shown for these time points. For T4 and T5, quantification of puncta distribution along the normalized proximodistal axis at 72h APF and adult are shown below the schematics (T4: adult quantification data same as Figure 3E; Rdl 72h APF, n = 4; GluClα 72h APF, n= 6. T5: adult quantification data same as Figure S4E, Rdl 72h APF, n = 7; GluClα 72h APF n= 2). Scale bars, 5 μm.

In sparsely labeled T4 and T5 neurons, Rdl accumulated as large puncta at the proximal region of dendrites at 48h APF, corresponding to the onset of dendrite extension (Figures 4A and 4B). As dendrites extend, Rdl remains enriched proximally with small fainter puncta distributed along their length. At 48 hrs, GluClα and nAChRβ1 puncta are also seen in incipient T4/T5 dendrites but over time disappear from the proximal region of the dendrite and accumulate more distally (Figures 4C-4D and S5E-S5F). This is particularly striking for GluClα. It accumulates throughout T4 and T5 dendrites at 48 and 72 hrs APF, disappears from T5 dendrites (consistent with the lack of glutamatergic inputs to T5) between 72 hrs APF and eclosion, and is retained only in the distal domain of T4 dendrites in the adult (Figures 4C and 4D).

The accumulation of NR subunits in developing L5 neurons was different. In contrast to T4, Rdl puncta were observed throughout L5 terminals in the medulla from 48 hrs through 72 hrs APF, but then disappeared from all but a highly restricted domain in medulla layers M1/M2 (Figure 4E). Also, in contrast to T4/T5 neurons, GluClα localized to terminals early, by 48 hrs APF, and this pattern remained into the adult (Figure 4F). Thus, different NR subunits localize to dendrites with distinct developmental dynamics and can be cell-type-specific rather than an intrinsic property of a given NR.

### A fly homolog of the synaptic adhesion protein ADAM22 forms a complex with GluClα

The exquisite localization of NR subunits to synapses in distinct dendritic domains raised the possibility that proteins associated with different NRs may regulate their localization, contribute to synaptic specificity, or both. As a first step to assessing this possibility, we focus on GluClα. Using an affinity purification mass-spectrometry (AP-MS) workflow, proteins associated with epitope tagged versions of GluClα were identified in extracts of fly brains. Two different tagged versions of GluClα and two complementary sets of controls were used (Figure 5A). Mind-meld (Mmd), a single pass transmembrane protein, was specifically bound with both versions of GluClα (Table S2). Co-IP experiments confirmed that Mmd interacts with GluClα in homogenates of brain tissue, but not with Rdl (Figure 5B).

**Figure 5.**
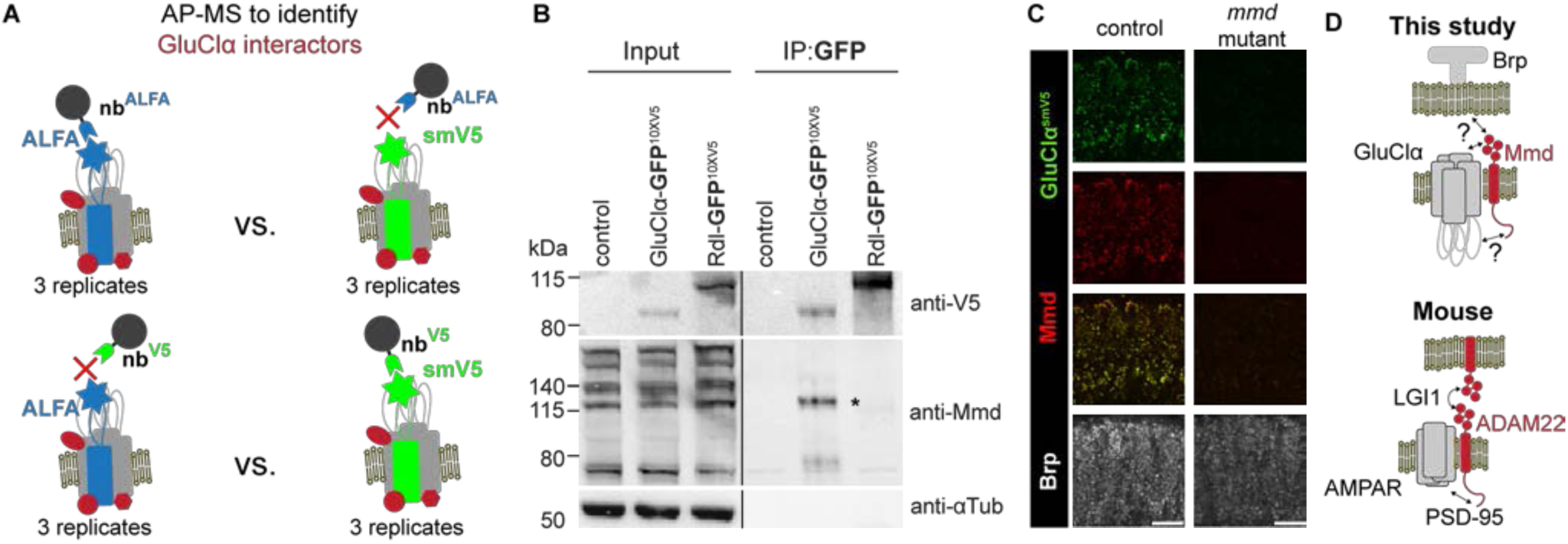
Mmd forms a complex with GluClα. (A) Design of AP-MS experiment used to identify interactors of GluClα. Tagged proteins were purified as indicated from head homogenates and subjected to mass spectrometry. Under these conditions, a single protein, Mmd, co-purified with both tagged versions of GluClα but not in controls (see Methods and Table S2). All samples represent 3 biological replicates. (B) Mmd with GluClα form a complex. Left panel, extracts from fly heads were probed for expression from flies carrying V5-tagged GluClα and V5-tagged Rdl with anti-V5 and anti-Mmd antibodies. Right panel, a nanobody against GFP was used to pull down GluClα or Rdl each tagged with smGFP-smV5. GluClα forms a complex with predominantly one form of Mmd. Asterisk denotes a weak band co-precipitating with tagged Rdl. (C) GluClα-smV5 staining is lost in an *mmd* mutant. Scale bar, 10 μm. (D) Bottom, structure of synaptic complex comprising ADAM22, the mouse homolog of Mmd. ADAM22 on the postsynaptic membrane and ADAM22 or a related protein on the presynaptic membrane are bridged by interactions with a dimer of LGI1 (Yamagata et al., 2018); Top, localization of GluClα and Mmd at a synapse in Drosophila as shown in this study. By contrast, interactions of Mmd with other proteins are not known.

As examined by confocal microscopy, there was complete overlap in the distribution of Mmd and GluClα (Figure 5C). To assess the role of Mmd in the formation of synapses containing GluClα, we analyzed brains mutant for *mmd*. Strong loss of mmd resulted in marked reduction in anti-GluClα staining (Figure 5C). Mmd is homologous to mammalian ADAM22 which has been shown to form a complex with AMPA receptors and acts as a synaptic adhesion molecule through its binding to LGI1 (Figure 5D) (Fukata et al., 2021). These data raise the possibility that in T4 neurons GluClα and Mmd form a complex selectively in the distal domain of their dendrites which, in turn, specifies the pattern of Mi9 inputs. We next sought to directly visualize GluClα and Mmd at these synapses.

### Mi9 axons form synapses juxtaposing GluClα and Mmd in distal T4 dendrites

EM studies revealed that seven to nine Mi9 axon terminals, organized into retinotopic columns, evenly distribute across the proximal, central, and distal domains of T4 dendrites (Figure 6A). Mi9 neurons synapse, however, only onto the distal-most domains (Figure 6A) (Shinomiya et al., 2019). Other neuron types form synapses within different domains along the proximodistal axis of these dendrites (Shinomiya et al., 2019). Mi9 is the only glutamatergic input to T4 dendrites. Here, we applied ExLSM to achieve sufficient resolution to visualize synapses between Mi9 and T4 dendrites and localize proteins within these synapses.

**Figure 6.**
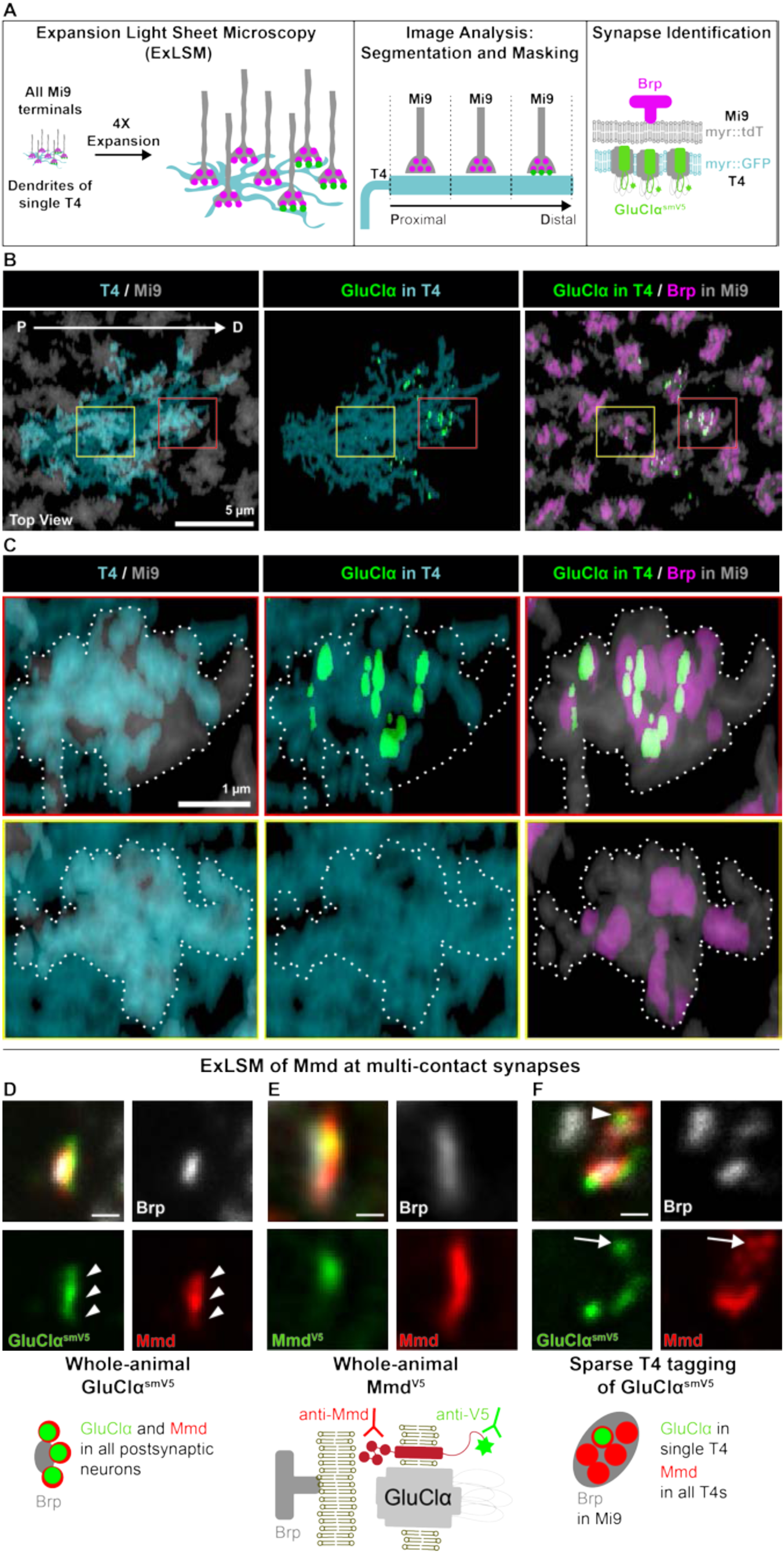
Mi9 axons form synapses juxtaposing GluClα and Mmd in distal T4 dendrites. (A) Identification of synapses between identified neuron types using ExLSM. Left panel, summary of sample preparation, imaging and synapse identification shown in panels B and C. Remaining panels, synaptic partners and pre- and postsynaptic proteins were visualized using four-channel light-sheet microscopy on expanded tissue (see Methods). Colors correspond to expression in Panels B and C. Arrow denotes proximo-distal axis along T4 dendrites. (B-C) Mi9 forms synapses selectively in the distal domain of T4 dendrites. B. T4 dendrites project across multiple columns. Mi9 axon terminals, one per column, contact all domains of T4 dendrites, but only form synapses distally. C. Upper row, synapses form between T4 and Mi9 in the distal domain (red box in B). Lower row, centrally located Mi9 terminals do not form synapses (yellow box in B). (D) GluClα co-localizes with Mmd at multi-contact synapses. Arrowheads point to multiple postsynaptic sites at a single multi-contact synapse co-labeled with Mmd and GluClα. (E) Mmd antibody recognizes an epitope in the Mmd ectodomain and the V5 antibody labels a V5 tag in the cytoplasmic tail of Mmd knocked into the genomic locus. (F) GluClα is tagged in a single T4 neuron. All other T4 synapses express Mmd but not tagged GluClα. Arrow points to a single postsynaptic site of a multiple-contact synapse co-labeled with Mmd and GluClα. Scale bars, 200 nm, represent unexpanded tissue size (adjusted for 4.65X expansion factor, see Methods).

To detect synapses between Mi9 and T4, we coupled labeling of GluClα in single T4 dendrites with staining of the membranes of Mi9 axon terminals and presynaptic Brp within them (Figures 6A and 6B). Control experiments verified our ability to assign Brp to individual Mi9 terminals and separate them from Brp in other processes in the neuropil (see Methods, Figures S6A and S6B). Consistent with EM studies, presynaptic sites in Mi9 terminals contacting the distal region of T4 dendrites were associated with GluClα expressed in these dendrites, while presynaptic sites of Mi9 terminals overlapping with the middle and proximal domains were not (Figures 6C and S6D-S6E). We next sought to determine whether Mmd also localized with GluClα at synapses between Mi9 and T4 dendrites.

We first confirmed via ExLSM the localization of Mmd and GluClα at synapses using both an antibody to the extracellular domain of Mmd and an antibody to an epitope tag inserted into its cytoplasmic domain (Figures 6D and 6E). To assess the localization of Mmd with GluClα in T4 neurons, we labeled single T4 neurons in which GluClα was selectively tagged with smV5 and co-stained with an anti-Mmd antibody. Each GluClα punctum co-localized with Mmd (Figure 6F).

Mmd staining extended beyond anti-GluClα immunoreactivity. This was expected, given that each Mi9 presynaptic site contacts multiple postsynaptic elements from other T4 neurons and only a single postsynaptic neuron was labeled with tagged GluClα at these synapses (Shinomiya et al., 2019). In summary, Mmd localized to the same synapses as GluClα in the distal domain of T4 dendrites juxtaposing Mi9 terminals.

## Discussion

In this study, we described a strategy to study single synapses between identified cell types by combining genetic tools, EM-based connectomics, and protein localization through expansion microscopy. To devise probes for specific synapses we developed tools to tag and map the distribution of endogenous NR subunits of receptors for the major neurotransmitters in *Drosophila* in single neurons and at single synapses between neurons. These studies highlighted the complexity of different NR subunit distributions in dendrites, the developmental dynamics of NR accumulation at different synapses, and their association with other proteins at subsets of synapses. These tagged receptors provide a resource for studies focusing on subcellular localization of neuronal proteins, the assembly of synapses and synaptic plasticity. As we discuss below, the unique features of wiring specificity of T4 and T5 dendrites suggest that in some developmental contexts targeting of NRs to discrete dendritic domains provides a spatial map of molecular determinants controlling synaptic specificity.

### Mapping NR subunits to specific synapses in single neurons

Our approach to conditionally tag receptors through modification of genomic loci preserves their endogenous mRNA expression levels, patterns, and temporal dynamics. We chose to insert epitope tags into poorly conserved and unstructured regions of variable length within a large cytoplasmic loop found in all cys-loop receptors. The distribution of the tagged NRs accurately reflects the localization of endogenous receptors. Tagged NRs selectively localized at sites adjacent to presynaptic partners (i.e., Brp and Cac) when visualized by either ExLLSM and ExLSM, and the distribution of receptors matched the neurotransmitter specificity of the neurons providing synaptic inputs. The increasing availability of genomes for comparative sequence analysis and the development of structure prediction tools such as AlphaFold (Jumper et al., 2021) facilitate the identification of sites to successfully tag other classes of NRs, as well as other synaptic proteins. The extensive synaptic connectivity maps in flies, reagents for manipulating specific cell types, and the array of endogenously tagged NR subunits reported here provides an opportunity to characterize the distribution of these receptors in many different circuits regulating a broad range of brain functions and behaviors.

The distribution of seven different tagged NR subunits for glutamate, GABA, and acetylcholine in the cys-loop superfamily in direction-sensitive T4 neurons was striking. The stereotyped patterns of each class of receptor matched the pattern of the neurotransmitters used by their respective presynaptic inputs. Unexpectedly, different NRs to the same neurotransmitter also localized to different domains receiving input from different presynaptic partners. This was seen for both cholinergic (excitatory) and GABAergic (inhibitory) synapses. The differential distribution of NR subunits to different spatial domains may contribute to the unique computational features of T4 dendrites (Strother et al., 2017). How these domains are established in neurons with diverse and often complex morphologies is not known. Perhaps the segregation of proteins to different domains may at a mechanistic level share features in common with the establishment of cell polarity domains in other cell types including apical basal polarity in epithelial cells (Buckley and Johnston, 2022).

Select cys-loop GABA receptors that differ in NR subunit composition are found in distinct domains in pyramidal cells of the mouse cortex (Contreras et al., 2019). It is likely that the complexity of receptor distributions in mammals also extends to different domains within the same dendrites as we have described here. In the mammalian brain there is a great diversity of cys-loop family of GABA receptors (i.e., GABAAR), with 19 distinct genes encoding GABAAR subunits (Contreras et al., 2019). Thus, tagging approaches in the mouse similar to what we report here in the fly may provide a way to uncover the spatial distributions of NR subunits and combinations of them at different synapses. As these receptors have different physiological properties, their spatial distributions may contribute to understanding information processing in dendrites of mammalian neurons.

### NR subunit diversity and synaptic specificity

Our analysis of T4 neurons raised the intriguing possibility that in some developmental contexts neurotransmitter receptors themselves may serve as recognition molecules specifying the pattern of pre-synaptic inputs. In these dendrites, each presynaptic neuron type forms synapses within restricted domains along the proximodistal axis (Takemura et al., 2017b) and these patterns correspond to the distribution of different NR subunits. Mi9 neurons, for instance, only form synapses within the distal domains of T4 dendrites while closely related Mi1 and Mi4 neurons form synapses in middle and proximal regions, respectively. This specificity is particularly striking as the dendritic arbors of many T4s overlap extensively such that each input axon terminal contacts the entire range of dendritic domains and yet makes synapses only within restricted spatial domains (Figure 7). A simple model to account for this specificity is that targeting of NR subunits to specific dendritic domains provides a spatial map of molecular signposts recognized by cell surface recognition proteins selectively expressed on the surface on different presynaptic terminal arbors. Alternatively, specific molecular determinants may be arranged along the proximodistal axis in discrete dendritic domains, and these may recruit both synaptic inputs and subsets of NR subunits to these sites. In either case, our studies suggest a close relationship between the targeting of NR subunits to different spatial domains and the specificity of synaptic inputs.

**Figure 7.**
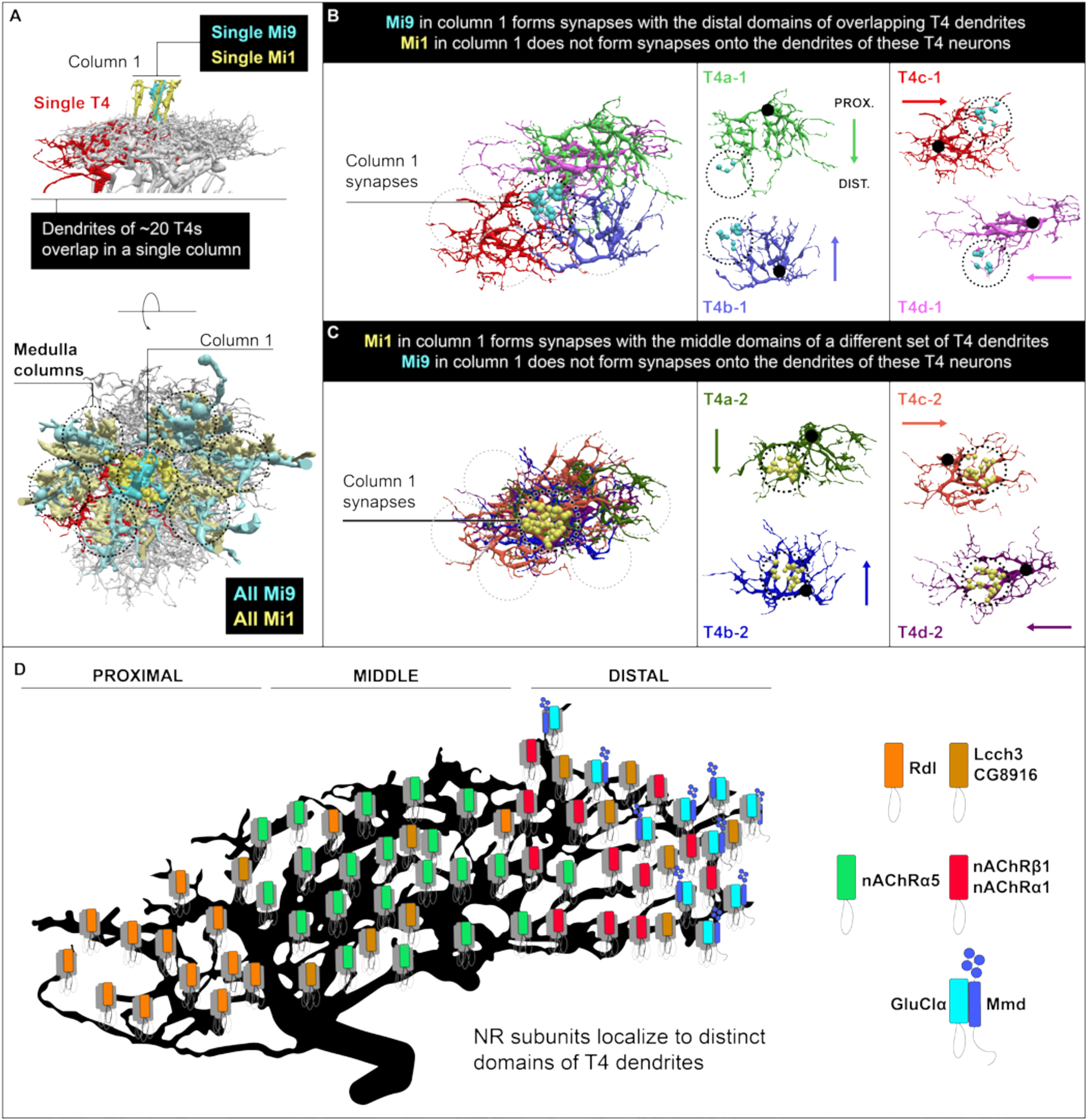
Model: NR subunits as synaptic specificity determinants in dendrites. Neurons providing presynaptic inputs into T4 dendrites in each column in the medulla are confronted by the full range of spatial domains due to the staggered overlap of many T4 dendrites from adjacent columns. These presynaptic inputs, however, discriminate between different T4 dendrites and are selective for specific spatial domains. Based on the connectome and our studies reported here, we propose that in some contexts NRs or proteins in complexes with them are sorted to specific spatial domains in dendrites and these act as synaptic specificity determinants. (A) Side and top views of 20 T4 dendrites contacting a single column. Upper panel (side view): The terminals of a single Mi9 and Mi1 in the central column (column 1) are depicted. Lower panel (top view): Dotted circular lines outline each medulla column. A single T4 dendrite is highlighted in red. Mi1 and Mi9 neurons in columns surrounding column 1 are shown in a slightly weaker shading than in column 1. The patterns of synaptic inputs to different T4s from Mi9 and Mi1 neurons in column 1 are shown in panels B and C. (B-C) All T4 dendrites receive inputs from both Mi9 and Mi1 neurons. In each column, different subsets of T4 dendrites receive input from a single Mi9 (B) or a single Mi1 (C) neuron. We describe one column as an example (column 1). Each circle demarcated with dotted lines indicates a different column; each column contains a single Mi9 and a single Mi1 axon terminal. Cyan indicates synapses established selectively by Mi9 in the distal domains of T4 dendrites (B), and yellow indicates synapses formed by Mi1 in the same column selectively with the central domains of a different subset of T4 dendrites (C). Synapses formed by Mi9 and Mi1 in the surrounding adjacent medulla columns are not shown. The right panels show the individual T4 neurons with distal domains overlapping (B) and overlapping middle domains (C). Colored arrows matching the dendrite colors indicate the proximal (PROX.) to distal (DIST.) polarity of these dendrites. The large black dots indicate the base of the dendrite in the proximal domain. (D) Summary illustrating the distribution of NR subunits across different domains of the T4 dendrites. The pattern of innervation of GABAergic, cholinergic, and glutamatergic input neurons matches the distribution of NR subunits (see Figure 3). NR subunits may serve as specificity determinants allowing presynaptic neurons to distinguish between different T4 dendritic domains. Alternatively, other spatial determinants, such as Mmd in T4, may recruit both NRs and presynaptic inputs to the same domain.

This model is consistent with studies in mammals. NRs have been shown to associate with other postsynaptic proteins that promote adhesion between pre, and postsynaptic membranes and these complexes may be selectively localized (Keable et al., 2020; Südhof, 2021). Here, for instance, we report the identification of a transmembrane protein, Mmd, in the distal domains of T4 dendrites in close association with GluClα receptor. A mammalian homolog of Mmd, Adam22, also co-localizes to synaptic glutamate receptors, albeit of a different class, and promotes adhesion between presynaptic and postsynaptic membranes (Figure 5D) (Fukata et al., 2021; Lovero et al., 2015; Yamagata et al., 2018). Alternatively, there is evidence that the extracellular domains of NR subunits can directly interact with proteins on the presynaptic membrane. For instance, the extracellular domain of the iGluR subunit GluA1 can interact with presynaptic neuronal pentraxin receptors, and this interaction can support the formation of synapses in a heterologous system (Lee et al., 2016). Similarly, the N terminal of the α1 GABAAR subunit has been shown to interact with neurexin-2β and modulate GABAAR function (Zhang et al., 2010). Together these observations raise the notion that, in some cases, the NR subunits themselves are intimately involved in matching of pre and postsynaptic membranes.

### Perspective

The complexity of neural circuit structure has become increasingly clear with the completion of dense EM connectomes (MICrONS Consortium, 2023; Scheffer et al., 2020; Shapson-Coe et al., 2021; Shinomiya et al., 2019; Takemura et al., 2013). Extensive studies have argued that different domains along the proximo-distal axis of T4 dendrites play a crucial role in direction-specific motion processing (Groschner et al., 2022; Gruntman et al., 2018; Strother et al., 2017).

The identification of different NRs and combinations of them within different domains provides an opportunity to understand computations at the molecular level.

How these specific molecular domains in dendrites form and their patterns of synaptic inputs emerge during development remains enigmatic. Localization of NR subunits to different domains at early stages of dendrite development raises the possibility that this molecular diversity contributes to determining the spatial distribution of specific synapses in dendrites. The use of tagged NRs and cell type specific manipulation with EM-based connectomics, ExLSM, genetics and biochemical methods provides a way of understanding how NRs become localized to specific domains and linking these to the specificity of synaptic inputs.

## Supporting information

Key Resource Table

## Acknowledgments

We thank members of the Zipursky laboratory for feedback on the manuscript. We thank M. DeSantis, D. Alchor of the Advanced Imaging Facility and K. Close, C. Christoforou, Y. He, A. Hu, P. Tillberg at the Janelia Research Campus for assistance and advice on acquisition and analysis of ExLLSM and confocal microscopy data. We thank Gokul Upadhyayula for discussion and advice on ExLSM image analysis. We thank Nathan Hwangbo of the Statistical Consulting Center of the UCLA Department of Statistics for assistance with statistical analysis. Reagents kindly provided by H. Hama, R. Davis, L. Luo, and M. Silies, as well as the Bloomington Drosophila Stock Center were critical for this work. Funding for this study in J.A.W laboratory was supported by the NIH (GM089778). This project was supported by an NIH BRAIN initiative grant (1RF1MH117823-01) in S.L.Z. laboratory and was supported by the Howard Hughes Medical Institute in Y.A. laboratory. S.L.Z. is an investigator of the Howard Hughes Medical Institute.

## Author Contributions (CRediT)

Conceptualization: P.S., S.L.Z.. Formal Analysis: Y.A., A.B., A.K., V.P., P.S., J.A.W.. Investigation: Y.A., H.B., A.B., P.G., A.K, Parmis, S.M., Pegah, S.M., V.P., P.S., J.Y., A.Y.. Methodology: Y.A., A.B., A.K., V.P., P.S., J.A.W.. Software: Y.A., A.K., V.P., P.S.. Supervision: Y.A., H.-S.L., J.A.W, S.L.Z.. Visualization: Y.A., A.B., A.K., P.S.. Writing – original draft: P.S., S.L.Z.. Writing – review & editing: A.K., P.S., S.L.Z.

## Declaration of Interests

The authors declare no competing interests.

**Table S1. List of tagged NR subunits and insertion sites**

Insertion sites relative to a reference protein isoform are reported, alongside inserted tags and gRNA target sequences.

**Table S2. Putative GluClα interactors as identified by AP-MS**

The statistical analysis package ArtMS was used to identify proteins that were differentially enriched during pull-down experiments using a nanobody against either the ALFA or V5 tag. Pull-downs followed by LC-MS/MS were conducted on triplicate samples of brain homogenates from either GluClα-1XALFA or GluClα-smV5 (Figure 5A) and proteins were quantified using MS1-based label free quantitation. Listed are proteins that showed significant enrichment in the experimental samples when compared to the negative control (e.g., nb ALFA on GluClα- 1XALFA vs nb ALFA on GluClα-smV5) and were identified using both a nanobody against the ALFA tag and the V5 tag. Imputed indicates proteins that were identified solely in the experimental sample and not in the negative control. For proteins that were not identified in the negative control, the measured fold-enrichment is undefined (#NUM!). In these cases, we imputed values in the negative control corresponding to the intensity of the least abundant proteins in that sample and then calculated the enrichment (iLog2FC) and p-values (iPvalue) after imputation.

## Supplemental figure titles and legends

**Figure S1. Tagged NR subunits localize to neuropils**

(A) Generation of constitutive and conditional alleles of epitope tagged NR subunits. A common knock-in construct for each NR subunit as shown in A was generated (Gratz et al., 2015; Kanca et al., 2019). Constitutive and conditional tagged receptors were generated as shown through passage of the knock-in through the germline carrying KD recombinase and Cre recombinase, respectively.

(B) Protein alignments of predicted NR subunit orthologs in multiple insect species (see Methods for details). Regions of low conservation within the cytoplasmic loop between transmembrane domains 3 and 4 were selected for tag insertion. Insertion sites are denoted by arrowheads and listed in Table S1.

(C) Expression pattern of tagged NR subunits visualized using antibodies targeting the epitope tag (anti-V5, anti-Ollas, anti-HA, or anti-ALFA). Schematic of the *Drosophila* optic lobe (La, lamina; Me, medulla; Lo, lobula; LP, lobula plate). Scale bar, 25 μm.

(D) Expression pattern of endogenous and tagged neurotransmitter receptor subunits as indicated using antibodies against the native protein (left) and the epitope tag (right). Scale bar, 25 μm.

(E) Brp with Cac at its center in medulla neuropil visualized using ExLLSM, as shown for the calyx in Figure 1H. Planar view is shown.

(F) nAChRβ1 subunits cluster juxtaposed to an active zone in the medulla.

(G and H) Examples of active zones and different NR subunits paired at synapses in the medulla (G) or calyx (H) neuropils.

(I) Example of Rdl clustered at active zones (arrows) as well as isolated puncta of Rdl not associated with Brp in the medulla (arrowheads). E-I scale bars, 200 nm.

**Figure S2. NR subunits localize at synapses of the mushroom body**

(A) A diagram of the MB circuit. Within the MB calyx, dendritic claw-like structures of multiple Kenyon cells (KCs) form synapses with the large axonal boutons of olfactory projection neurons (PN). In contrast, thin axons of KCs form converging synapses onto the dendrites of individual mushroom body output neurons (MBONs) in specific compartments of the mushroom body lobes (Takemura et al., 2017a).

(B) nAChRα6-smOllas in Kenyon cells. Inserts show magnified views of the calyx where olfactory PNs form diverging synapses with dendrites of KCs.

(C) Rdl-smV5 (bottom) and myr-tdTomato (top) in glutamatergic MBON05 which arborize dendrites in the γ4 compartment and send axonal projections in the γ1 and γ2 compartments and outside the MB.

(D) Rdl-smV5 (bottom) and myr-tdTomato (top) in GABAergic MBON11 that arborize dendrites in the core of distal peduncle and γ1 compartment (γ1pedc) and send axonal projections to the α/β lobes and outside the MB.

(E) ExLLSM images of the constitutively tagged nAChRα6-smOllas and Brp labeled by nc82 antibody in the MB calyx. Projection of 6 frames is shown.

(F) Magnified view of the inset shown in E highlighting a stereotypical microglomerulus where a large axonal bouton of a putative PN is surrounded by dendritic claws of KCs. Signal is organized in accordance with the characteristic arrangement of synaptic structures, as seen by EM.

(G) nAChRα6-smOllas and Brp in the α3 compartment of the MB lobe.

(H) A magnified view of G. The arrow indicates a putative “rosette synapse” where multiple KCs converge onto a single dendrite within the dendritic arbor of an MBON. The arrowheads indicate putative rosette synapses that do not stain for nAChRα6-smOllas.

(I-L) ExLLSM images of the constitutively tagged nAChRα1-smOllas and Brp in the calyx (I-J) and the α3 compartment (K-L).

(M) Colocalization analysis, as summarized in N-P, for nAChRα6-smOllas in the calyx. Top, distribution of distances between nAChRα6-smOllas and Brp for colocalized receptors (orange) and for receptors that are not colocalized with Brp (blue) (see Methods). Bottom left, the size of nAChRα6-smOllas objects plotted against distance to the nearest Brp. Bottom right, the frequency distribution of size of nAChRα6-smOllas objects. nAChRα6-smOllas objects were colocalized with Brp if they had at least a single voxel overlap or contact (see Methods).

(N) The percentages of tagged-receptor signals (i.e., voxels above the threshold) in Brp- colocalized objects in the original (orange bars) and rotated (gray bars) images. Dots indicate means of each 360x480x480 voxels tile. 8-24 tiles were analyzed for each condition. The left most “nc82” bars are control experiments in which Brp was detected with nc82 primary antibody and two different secondary antibodies. All tagged-receptor signals were significantly enriched in Brp-colocalized objects compared to rotated images, except for Lcch3-smOllas in the MB α3 compartment. Sidak’s multiple comparison test was used for the analysis.

(O) The numbers of tagged-receptor objects that colocalized with Brp (orange bars) or that did not colocalize (blue bars).

(P) The sizes of tagged-receptor objects that colocalized with Brp (orange bars) or that did not colocalize (blue bars). Brp-colocalized receptor objects are significantly larger than ones that did not colocalize with Brp, except for Lcch3-smOllas in the calyx and the MB α3 compartment, and nAChRα7 in the MB α3 compartment. Sidak’s multiple comparison test was used for this analysis.

Scale bars in B and E-L panels represent unexpanded tissue size (adjusted for 4.65X expansion factor, see Methods).

**Figure S3. Tagged neurotransmitter receptor subunit distributions in Tm3 dendrites**

(A) NR subunit mRNA expression in optic lobe neurons from scRNA-seq at 96h after pupal formation (Kurmangaliyev et al., 2020). Bold type, tagged alleles generated in this study.

(B) Detailed step-by-step schematic illustrating the recombinase cascade that triggers the conditional tagging of NR subunits in sparsely distributed neurons.

(C-E) Localization of GluClα-smV5 (C), nAChRb1-smHA (D) and Rdl-smV5 (E) in Tm3 dendrites of the medulla. Left, morphology of reconstruction of the EM-based inputs to Tm3 neurons and annotated synapses for each neurotransmitter type; question mark on magenta synapses represents inputs from neurons of unknown neurotransmitter specificity (Takemura et al., 2015). Right panels, neuron morphology (gray) and NR subunits in color as indicated. Scale bars, 5 μm.

(F) Quantification of data shown in A-C for several neurons (EM n=3, Rdl n=6, GluClα n=9, nAChRβ1 n=13) within each layer and of three neurons from the EM data. Bonferroni adjusted p-value (*) < 0.05 from Wilcoxon rank-sum test.

**Figure S4. Localization of NR subunits on T4 and T5 dendrites**

(A) Upper panel, EM reconstruction of different presynaptic inputs, as indicated, along the proximodistal axis of T5 dendrites. Dendrites span an average of three columns. Lower panel, annotated synapses for different neurotransmitter inputs are shown. Colored arrows show domains targeted by color matched presynaptic inputs from the upper panel. TmY15 is an exception forming synapses across the length of T5 dendrites.

(B-D) Analysis of NR subunit localization in T5 dendrites as in Figures 3B-3D. Scale bars, 5 μm.

(E) Quantification of NR subunit expression along the normalized proximo-distal axis of T5 dendrites (Rdl, n = 7; Lcch3, n = 3; CG8916, n = 2; GluClα, n = 6; nAChRα1, n = 8; nAChRα5, n = 7; and nAChRβ1, n = 10).

(F) Quantification of inputs to T4 and T5 dendrites from EM reconstructions (upper panel) and this study (lower panel). Quantification of puncta from Figures 3B-3D and Figures S4B-S4D. T4 EM, n = 20. T4 immunofluorescence: Rdl, n = 11; CG8916, n = 6; Lcch3, n = 8; GluClα, n = 8; nAChRα1, n = 7; nAChRα5, n = 6; and nAChRβ1, n = 28. T5 EM, n = 20; T5 immunofluorescence: Rdl, n = 13; CG8916, n = 4; Lcch3, n = 4; GluClα, n = 6; nAChRα1, n = 12; nAChRα5, n = 8, and nAChRβ1, n = 19. Bonferroni adjusted p-value (***) < 0.001 from Wilcoxon rank-sum test for EM; p-value (***) < 0.001 from Wilcoxon rank-sum test for immunofluorescence.

(G) Genetic scheme for labeling two different NR subunits in the same neuron. The approach is analogous to the one described in Figure S3B (see steps 1-3), with the addition of an allele for the second subunit tagged with a different epitope.

(H and I) Double labeling of nAChRβ1-smHA with either nAChRα5-smOllas or nAChRα1- smOllas in single T5 dendrites (H) or T4 dendrites (I). (H) Within the overlapping region (vertical dashed lines), nAChRβ1-smHA and nAChRα5-smOllas puncta are separate, indicating that these are not at the same synapses. By contrast, nAChRβ1-smHA and nAChRα1-smOllas localization domains within T5 dendrites have nearly complete overlap. Some puncta appear co- localized, however at this level of resolution it is not clear whether these NR subunits localize to the same synapses. (I) Double labeling of nAChRβ1-smHA with either nAChRα5-smOllas or nAChRα1-smOllas in single T4 dendrites. Scale bars, 5 μm.

**Figure S5. Developmental dynamics of NR subunits expression**

(A) Developmental expression dynamics of mRNAs of Rdl, nAChRβ1 and GluClα in distinct neuron types of the developing optic lobe as derived by scSeq data analysis (Kurmangaliyev et al., 2020). T4, T5 and L5 expression are colored as indicated.

(B-D) Immunofluorescence staining of NR subunits during development in the medulla neuropil of the optic lobe and neuronal cell bodies present in the above cortical layer. Dashed lines denote the boundary that separates the cell body layer located above the synapse-rich neuropil of the medulla. NR subunits probed in B-D, as indicated.

(E and F) Time course of nAChRβ1-smHA subunit accumulation during development of T4 (E) and T5 dendrites (F). Times shown are 48h APF, 72h APF and adult (Ad). Schematic of dendrite development with NR subunits distribution along the normalized proximodistal axis at 72h APF and adult are shown. (T4: Adult same as in Figure 3E, nAChRβ1 72h APF n = 10: T5: Adult same as in Figure S4E, nAChRβ1 72h APF n=10).

**Figure S6. Strategy and validation of labeling of single synapses using ExLSM**

(A-B) (A) Brp-smV5 was selectively tagged in Mi9 neurons using synaptic tagging after recombination (STaR) (Chen et al., 2014b), in combination with native Brp staining using the monoclonal anti-Brp antibody nc82. (B) Six examples of Mi9 axon terminals are shown with conditionally tagged Brp-smV5 and extracted nc82 signal in labeled Mi9 neurons. A separate analysis pipeline was developed for 4X-expansion light-sheet microscopy data acquired using a commercial light-sheet microscope (Zeiss LS7) (see Methods).

(C-E) (C) Genetic strategy used to label GluClα in single T4 dendrites and Mi9 terminals. Presynaptic sites were labeled with Brp, and the signal was segmented and assigned to Mi9 terminals (see Methods). (D-E) Representative examples from additional replicates in independent animals as in Figures 6B-6C.

## STAR Methods

### RESOURCE AVAILABILITY

#### Lead contact

All requests for additional information and reagents should be directed to and will be fulfilled by the lead contact, S. Lawrence Zipursky.

#### Materials availability

Flies generated in this study have been deposited to the Bloomington Drosophila Stock Center. Plasmids have either been deposited to Addgene or are available upon request.

#### Data and Code Availability

Any additional information required to reanalyze the data reported in this paper is available from the lead contact upon request.

### EXPERIMENTAL MODEL AND SUBJECT DETAILS

#### Fly husbandry

*Drosophila melanogaster* was reared on cornmeal/molasses medium at 25°C in a humidity-controlled incubator. Females were dissected for experiments unless otherwise noted. White prepupae were collected for developmental studies and designated as 0h after pupal formation (APF). Stocks used and generated in this study are listed in the key resources table. Genotypes used in each figure panel and related immunofluorescence staining conditions are reported in Table S3. Transgenic flies were generated by integrating DNA constructs into specific landing sites, while targeted alleles were created through CRISPR-mediated homologous recombination as described below. This was carried out using a commercial injection service (BestGene, Inc.).

### METHOD DETAILS

#### Identification of NR subunit genes

NR subunit gene numbers were obtained from https://www.genenames.org (human), https://www.informatics.jax.org (mouse) or https://www.flybase.org (fly). For each organism, Cys-loop NR subunits, iGluR subunits and GPCR neurotransmitter receptor subunit were compiled. GPCRs responding to neuropeptides were omitted from the counts.

#### Selection of tag insertion sites

All tags were inserted within the unstructured intracellular loop between the third (M3) and fourth (M4) transmembrane domains. In addition, we chose insertion sites that were poorly conserved and favored ones with evidence of sequence insertion in other species. We generated sequence alignments with the Clustal Omega program in uniprot (https://www.uniprot.org/align). Closely related (e.g., *D. yakuba*) and distantly related (e.g., *T. castaneum*) insect species were used for alignment. The insertion sites are indicated in Figure S1 and summarized in Table S1. Species abbreviations - DROME: *Drosophila melanogaster;* DROSE: *Drosophila sechellia;* DROSI: *Drosophila simulans;* DROYA: *Drosophila yakuba;* DROAN: *Drosophila ananassae;* DROPS: *Drosophila pseudoobscura;* DROPE: *Drosophila persimilis;* DROWI: *Drosophila willistoni;* DROMO: *Drosophila mojavensis;* DROGR: *Drosophila grimshawi;* AEDAE: *Aedes aegypti;* ANOGA: *Anopheles gambiae;* CULSO: *Culicoides sonorensis;* TRICA: *Tribolium castaneum*.

#### Molecular biology

##### Generation of conditional tag cassettes

We generated a conditional tag cassette, pBS-KDRT-STOP-loxP-3XP3::dsRed-loxP-STOP-KDRT-smGFPTag, for the tags smGdP-10XV5, smGdP-10XHA, and smGdP-10XOllas. To generate the cassette, we used a combination of restriction enzyme-based cloning and HiFi DNA Assembly (NEB cat #E2621). First, we replaced the sequence between MluI and MfeI in KDRT-STOP-STOP-KDRT from pJFRC164 (Addgene plasmid #32141) with loxP-3XP3::DsRed-loxP from pHD-DsRed-attP-w+ (Addgene plasmid #80898). The 3xP3::DsRed marker allowed easy screening for successful genomic insertions which are subsequently removed by germline expression of Cre recombinase. Next, we added a *Drosophila* codon-matched GS linker and the coding sequence for each tag downstream of KDRT. We cloned the sequence for each tag from the following plasmids: pJFRC206 (Addgene plasmid #63168) for smGdP-10XV5, pJFRC201 (Addgene plasmid #63166) for smGdP-10XHA, and pJFRC210 (Addgene plasmid #63170) for smGdP-10XOllas. To place the first KDRT sequence in frame, the cassette was preceded by the dinucleotide GG to encode a glycine with the first nucleotide of the KDRT sequence, and then the entire cassette was cloned into pBlueScriptII KS(-) (Agilent Cat# 212208) between PciI and XbaI. A second PciI site was inserted upstream of XbaI to facilitate cassette linearization in subsequent steps. All plasmids were sequence validated by Sanger sequencing. smGdPTag encodes a GFP protein with 10X epitope tags distributed amongst the C and N termini, and one of the loops (Viswanathan et al., 2015). In addition to the inserted tags, the GFP sequence of smGdP contains amino acid substitutions that render GFP non-fluorescent (GdP: fluorescence dead GFP – (Nern et al., 2015). For simplicity we refer to smGdPTag throughout the study as smGFP-Tag or smTag.

We generated a conditional 1XALFA cassette by replacing smGdP-X from pBS-KDRT-STOP-loxP-3XP3::dsRed-loxP-STOP-KDRT-smGFP-Tag with Drosophila codon-optimized 1XALFA followed by a GS linker. ALFA-tag is a commercially developed epitope tag which forms a small and stable α-helix and is recognized by a high affinity nanobody (Götzke et al., 2019). Detailed protocols are available upon request. The plasmids and sequences have been deposited in Addgene.

##### Generation of pU6-gRNA

We identified gRNA target sequences that cut within 1-11 nt of the selected insertion site with an efficiency score above 5, as defined by the CRISPR Efficiency Predictor (https://www.flyrnai.org/evaluateCrispr/). The gRNA sequence oligos were synthesized (Integrated DNA technologies) with the forward oligo having a TTCG overhang at the 5’ end and the reverse oligo having an AAAC overhang added to the 5’ end for subsequent ligation into pU6. After annealing, the oligos were ligated into BbsI-linearized pU6b-sgRNA-short (Ren et al., 2013). All pU6 vectors generated were verified by Sanger sequencing. The gRNA sequences used in this study are listed in Table S1. gRNA1, used for generating alleles using short homology arms (see below – (Kanca et al., 2019), was cloned into pU6 as described for gene specific gRNAs.

##### Generation of donor constructs

The generation of donor constructs involved two different methods for different sets of alleles. For the first set, long homology arms of approximately 1kb were used for homologous recombination of the conditional tag cassette. To improve cloning efficiency as well as donor integration by homologous recombination, we took advantage of a second method which employs shorter homology arms of around 100bp in combination to in vivo directed linearization of the donor vector, as described (Kanca et al., 2019). This strategy was used to generate alleles of Grd, CG8916 and nAChRα3, as well as the 1XALFA tagged allele of GluClα.

For the generation of alleles using long homology arms, we employed HiFi DNA assembly (NEB Cat# E2621) to assemble the donor constructs. The long homology arms (∼1kb) were PCR amplified and inserted into pHD-DsRed-attP-w+ (Addgene plasmid #80898), which was linearized with XhoI and EcoRI. The tag cassette was introduced by cloning in the PciI linearized conditional tag cassette of choice. For smGFP tags, we included a GS linker in the primer used to generate the 3’ homology arm. In contrast, the 1XALFA tag contains the GS linker within the conditional cassette. All pHD-geneX donor plasmids were sequence validated. Single nucleotide polymorphisms (SNPs) were permitted in intronic regions and in coding regions only when leading to synonymous codon substitutions from the dm6 reference genome.

For the generation of alleles using short homology arms, we utilized HiFi DNA assembly (NEB) with two steps. In the first step, we synthesized the donor homology arms into pUC57-Kan (Genewiz, Inc.), with each of the two ∼125bp homology arms flanked on the outside by gRNA1 target and PAM sequences and DNA assembly-specific homology arms matched to the chosen PciI linearized conditional cassette. The two arms were separated by a random sequence linker flanked by restriction sites that were absent in the homology arms. We included a GS linker in the synthesized sequence for the generation of donors with smGFP tags, while the 1XALFA conditional cassette included the GS linker within the cassette. In the second stage, we linearized pUC57-Kan-geneX with restriction enzymes recognizing the sites within the linker and cloned in the PciI linearized fragment that encoded the required conditional cassette using HiFi DNA assembly (NEB Cat# E2621). All donor vectors were confirmed by Sanger sequencing. Plasmids and sequences are available upon request.

##### Generation of pJFRC-10XUAS-FRT-STOP-FRT-myrFP-2A-KDR::Pest

HIFI DNA assembly (NEB Cat# E2621) was used to generate either 10XUAS-FRT-STOP-FRT-myr::GFP-2A-KDR::PEST or 10XUAS-FRT-STOP-FRT-myr::tdTomato-2A-KDR::PEST. The GFP coding sequence of pJFRC177 was replaced either by GFP-2A (cassette C: GS linker- FRT-STOP-FRT-GFP-2A-LexAVP16, (Chen et al., 2014b) or tdTomato-2A (UAS-DIPalpha-2A- tdTomato) (Xu et al., 2018), both followed by the coding sequence of KDR::PEST recombinase from pJFRC161 (Addgene: 20XUAS-IVS-KD::PEST plasmid #32140, Nern et al., 2011). Plasmids and sequences are available upon request.

##### Generation of pJFRC-13XlexAoP-10XUAS-FRT-STOP-FRT-myrFP-2A-KDR::Pest

A combination of HIFI DNA assembly (NEB) and restriction enzyme-based cloning was used to generate 13XLexAoP2-FRT-STOP-FRT-myr::GFP-2A-KDR::PEST through modification of pJFRC177 (Addgene: 10XUAS-FRT-STOP-FRT-myrGFP, plasmid #32149, Nern et al., 2011). First, the 10XUAS sequence of pJFRC177 was replaced by 13XLexAoP2 from pJFRC19 (addgene:13XLexAoP2-IVS-myrGFP, plasmid #26224, Pfeiffer et al., 2010). Second, the GFP coding sequence of pJFRC177 was replaced either by GFP-2A (cassette C: GS linker-FRT-STOP-FRT-GFP-2A-LexAVP16, (Chen et al., 2014b) followed by the coding sequence of KDR::PEST recombinase from pJFRC161 (Addgene: 20XUAS-IVS-KD::PEST plasmid #32140, Nern et al., 2011). Plasmids and sequences are available upon request.

#### Generation of tagged NR subunit alleles

For alleles generated using donor vectors with long homology arms, pHD-geneX donor plasmid and pU6-geneX-gRNA were injected into flies expressing Cas9 in the germline by BestGene Inc. Successful integration of the donor cassette was identified through expression of DsRed in the eyes and negative for expression of mini-white in the eye. Resulting flies were PCR validated for correct insertion within the selected insertion site (Table S1). To generate the conditional tagged allele, we crossed flies to a line expressing Cre recombinase (BDSC# 1092) to excise loxP flanked DsRed from the STOP cassette within the targeted allele (Figure S1A). To generate the whole fly constitutively tagged allele, we crossed the conditional tagged allele to a line expressing KD recombinase in the germline (w; 3XUAS-KDR; nos-gal4::VP16) to excise the KDRT flanked STOP cassette upstream of the tag within the targeted allele. Resulting conditional and whole fly constitutively tagged alleles were balanced. When the whole fly constitutively tagged allele was not homozygous viable, we backcrossed flies three times with w1118 to clean out any off target CRISPR events that could underlie lethality. We were not able to outcross homozygous lethality for Rdl-KDRT-smV5, Rdl-KDRT-smHA and GluClα- KDRT-smV5. To test if lethality was due to insertion of the tag within the NR subunit locus or to off target CRISPR induced mutations, we tested viability over deficiency lines (Rdl Df: BDSC# 8066, GluClα Df: BDSC# 8964) or loss of function alleles for Rdl (Rdl^1^, BDSC#1687). As these animals were viable, lethality in homozygous animals does not reflect disruption of receptor function due to the tag. All tagged alleles were sequence validated to confirm correct cassette excision. SNPs were allowed in intronic regions and in coding regions only when resulting in synonymous codon substitutions.

When generating alleles using short homology arms, pUC57-geneX donor plasmid, pU6-geneX- gRNA and pU6-gRNA1 were injected into flies expressing Cas9 in the germline by BestGene Inc. Generation of flies was carried out as described above with the exception that flies were only screened for expression of DsRed in the eye. Conditionally tagged and whole fly tagged alleles of NR subunits were generated as described above.

#### Tagging of NR subunits in single neurons of the optic lobe

Sparse labeling of cells was achieved as outlined in Figure S3B. To achieve sparse labeling of cells, we optimized timing of heat shock to mediate Flp-out of the FRT-STOP-FRT (FSF) cassette from 10XUAS-FSF-myrFP-2A-KDR::Pest. To achieve sparse labeling of neurons found in each column or more than one per column, we used a less efficient variant of hsFlp (hsFlpG5::Pest(Opt) BDSC#77140). 0-24h APF pupae carrying hsFlp, 10XUAS-FSF-myrFP-2A- KDR::Pest, the cell type specific GAL4 and the endogenous conditionally tagged allele of the NR subunit of interest (or more than one allele if more than one receptor was investigated, see Figures S4G-S4I), were heat-shocked at 37°C (Tm3:15 min, T4T5: 8-12 min, L5: 8 min) and subsequently reared at 25°C. The degree of labeled cells is very sensitive to small changes in time and temperature. The conditions for labeling were empirically established for every GAL4 driver. Brains for analysis were dissected from 1-5 day old flies. For developmental studies of NR subunit expression in T4/T5 and L5 neurons, the same procedure was followed as above, and brains were dissected at either 48h APF or 72h APF.

#### Immunostaining

For adult fly brains, 1-5 day old flies were decapitated, and the brain was dissected in ice cold Schneider Medium (SM) (Thermo Scientific, Cat# 21720001). Up to three brains were kept on ice in SM prior to fixation in a single well of a Terasaki plate (Thermo Scientific, Cat# 163118). All subsequent steps were carried out in Terasaki plates. Brains were fixed overnight at 4°C in glyoxal acid-free fixative (Addax Biosciences, Cat# VI25) supplemented with 5% (w/v) sucrose (addaxS) or in a fixative containing 3% glyoxal at pH 5.0 (3% glyoxal fixative) (Richter et al., 2018), as indicated in Table S3. When fixed with 3% glyoxal fixative, brains were fixed for 30 min at room temperature, followed by 30 min quenching in 0.1 M NH_4_Cl. Fixation in acid-free glyoxal with 5% sucrose improves preservation of fine neuronal morphology compared to 3% glyoxal fixative. Both fixatives improve immunostaining of NR subunits and Bruchpilot, compared to standard 4% PFA in PBS. After fixation, brains were washed 3X and incubated for 2 hours at RT in PBSTX (PBS with 0.5% (v/v) Triton X-100 (Sigma Aldrich Cat# T8787) with addition of 10% normal goat serum (PBSTN) (Sigma-Aldrich Cat# G6767). Brains were incubated for 2-3 days at 4°C with primary antibody mix in PBSTN, subsequently washed 2X 2hrs with PBSTX and further incubated 2-3 days overnight with secondary antibodies in PBSTN. Secondary antibody was washed out by incubating brains 2X for 2 hrs in PBSTX and subsequently mounted as described below.

The following primary antibodies and concentrations were used for samples imaged using confocal microscopy. Rabbit anti-nAChRα6 (1:1,000; Nakayama et al., 2016), rabbit anti- nAChRα7 (1:1,000, courtesy of H. Hama), rabbit anti-Rdl (1:100; Liu et al., 2007), rabbit anti- GluClα (raised against a peptide from *Musca domestica* GluCl, which is conserved in D. melanogaster, 1:500, Kita et al., 2013), rabbit anti-Mmd (1:500 Guo et al., manuscript under preparation), rabbit anti-Ollas (1:10,000; GenScript, Cat# A01658), rat anti-Ollas (L2, Novus Cat# NBP1-06713, RRID:AB_1625979), rat anti-HA (3F10, Roche Cat# 11867423001, RRID:AB_390918), mouse IgG2a anti-V5 (SV5-Pk1, 1:500; Abcam Cat# ab27671, RRID:AB_471093), chicken anti-V5 (1:500, Abcam Cat# ab9113, RRID:AB_307022), mouse anti-V5-Tag:DyLight®550 (Bio-Rad Cat# MCA1360D550GA, RRID:AB_2687576), FluoTag X2 anti-ALFA::Atto488 (1:500; NanoTag Biotechnologies, Cat# N1502-At488), FluoTag X4 anti-RFP::AZDye568 (1:500; NanoTag Biotechnologies, Cat# N0404-AF568), rabbit anti-DsRed (1:200, Takara Bio Cat# 632496, RRID:AB_10013483), guinea pig anti-RFP (1:1,000, Synaptic Systems, Cat# 390004, RRID:AB_2737052) and chicken anti-GFP (1:1000, Abcam Cat# ab13970, RRID:AB_300798). Secondary antibodies were used at 1:500 for confocal microscopy. From Abcam: Alexa Fluor 488 goat-anti-rat (Cat# ab150165, RRID:AB_2650997), Alexa Fluor 568 goat-anti-rat (Cat# ab175710, RRID:AB_2832918), and Alexa Fluor 488 goat-anti-rabbit (Cat# ab150081, RRID:AB_2734747). From Biotium: CF405S goat-anti-mouse IgG1 (Cat # 20380), CF633 goat-anti-rabbit (Cat# 20123-1, RRID:AB_10853138) and CF633 goat-anti-mouse (Cat# 20341). From Invitrogen: Alexa Fluor 488 goat-anti-mouse (Cat# A-11029, RRID:AB_2534088), Alexa Fluor 568 goat-anti-mouse IgG1 (Cat# A-21124, RRID:AB_2535766), Alexa Fluor 488 goat-anti-mouse IgG2a (Cat# A-21131, RRID:AB_2535771), Alexa Fluor 546 goat-anti-mouse IgG2a (Cat# A-21133, RRID:AB_2535772), Alexa Fluor 568 goat-anti-guinea pig (Cat# A-11075, RRID:AB_2534119), Alexa Fluor 488 goat-anti-rabbit (Cat# A-11034, RRID:AB_2576217), Alexa Fluor 568 goat-anti-rabbit (Cat# A-11036, RRID:AB_10563566), Alexa Fluor 647 goat-anti-rabbit (Cat# A-21244, RRID:AB_2535812), Alexa Fluor Plus 488 goat-anti-chicken (Cat# A32931, RRID:AB_2762843), and Alexa Fluor 488 goat-anti-chicken (Cat# A32931, RRID:AB_2762843). From NanoTag Biotechnologies: Atto 647N FluoTag X2 anti-mouse IgG1 (Cat# N2002- Atto647N) and AbberiorStar635P FluoTag X4 anti-rabbit (Cat# N2404-Ab635P). From Jackson ImmunoResearch Labs: Alexa Fluor 488 goat-anti-rat (Cat# 112-545-167, RRID:AB_2338362), Alexa Fluor 488 goat-anti-chicken (Cat# 103-545-155, RRID:AB_2337390) and Alexa Fluor 488 goat-anti-mouse (Cat# 115-545-166, RRID:AB_2338852).

#### Preparation of samples for NR subunit tagging in cell types of the mushroom body

Single neurons of the mushroom body were generated as detailed above and outlined in Table S3. A 2 hr heat shock at 24-48 hrs APF was chosen to generate single neuron labeling.

Dissection and immunohistochemistry of fly brains were carried out as previously described with 3% Glyoxal fixative instead of 2% PFA (Jenett et al., 2012) using the antibodies listed in Table S3. Brains of 5-day-old female flies were dissected in Schneider’s insect medium and fixed in 3% Glyoxal fixative (3% v/v glyoxal (Sigma-Aldrich, Cat# 128465), 5% v/v ethanol, 0.75% v/v acetic acid, pH 5.0) for 2 hr at room temperature (RT). The samples were quenched in 0.1M NH_4_Cl for 30 min followed by four washes in PBT (0.5% Triton X-100 in 1x-PBS), 10 min each. After washing in PBT, tissues were blocked in 5% normal goat serum (or normal donkey serum, depending on the secondary antibody) for 90 min. Subsequently, tissues were incubated in primary antibodies diluted in 5% serum in PBT for 2 days on a nutator at 4°C, washed four times in PBT for 30 min, then incubated in secondary antibodies diluted in 5% serum in PBT for 2 days on a nutator at 4°C. Tissues were washed thoroughly in PBT four times for 30 min or longer.

#### Sample mounting for confocal microscopy

Samples were mounted using either DPX mountant (EMS, Cat# 13510) as previously described (Nern et al., 2015) or Everbrite mounting medium (Biotium, Cat# 23001), as denoted in Table S3. DPX facilitates imaging of neurons deep in the brain. The refractive index of DPX matches the refractive index of the immersion oil used with high NA objectives, facilitating higher resolution achievable using Airyscan.

#### Confocal microscopy

All confocal images were acquired on a Zeiss LSM880 with 405 nm, 488 nm, 561 nm, and 633 nm lasers. Images were acquired with a Plan-Apochromat 63x/1.4 Oil DIC M27 objective for single cell and whole brain imaging or a Plan-Apochromat 40xx/1.2 Imm Korr DIC M27 for whole optic lobe imaging (Figure S1).

Airyscan was used in conjunction with 63X objective imaging in RS mode to maximize signal capture and increase resolution compared to conventional confocal imaging. All samples were imaged using precision cover glasses with #1.5H thickness (Thorlabs, Cat# CG15CH2) and the Airyscan detector alignment was verified and adjusted prior to imaging each sample. Frame size and Z step was optimal to achieve maximal resolution. Images were processed using Zen Blue 2.3 with Airyscan processing set to auto-filter and 3D processing.

#### ExLLSM sample preparation and imaging

The samples for Expansion Microscopy (ExM) were dissected, fixed and immunostained as stated for neurons of the mushroom body and staining was done sequentially to avoid cross-reactivity. All samples were processed using a protein retention ExM protocol with minor modifications (Asano et al., 2018; Tillberg et al., 2016). All solutions were prepared in milliQ-grade water unless otherwise stated: AcX stock (acryloyl-X, SE ThermoFisher, A20770) at 10 mg/mL in anhydrous DMSO (Sigma, Cat # 276855); PLL solution (Ted Pella, Cat# 18026) with Photo-Flo detergent (EMS, Cat# 74257) added 1:500 (v/v); Acrylate stock at 4M, prepared by neutralizing 5.5 mL acrylic acid (99% purity; Sigma, Cat# 147230) with 10N NaOH using a water bath and fume hood, in a total volume of 20mL (Tillberg, 2021); Acrylamide stock at 50% (w/v) (Sigma, A9099); and Bis-acrylamide stock at 1% (w/v) (Sigma, M7279). Monomer stock: 11.5 mL sodium acrylate stock, 2.5 mL acrylamide stock, 7.5 mL bis-acrylamide stock, 18 mL 5 M NaCl (Sigma, Cat# S5150-1L), 5 mL 10xPBS (ThermoFisher, Cat# 70011044), and 2.5 mL water for a total volume of 47 mL. 4HT stock: 4-hydroxy-TEMPO at 0.5% (w/v) (Sigma, Cat# 176141). TEMED stock: N,N,N′,N′-Tetramethylethylenediamine at 10% (v/v) (Sigma, Cat# T7024). APS stock: ammonium persulfate at 10% (w/v) (Sigma, Cat# A3678). ProK digestion buffer: 0.5% Triton X-100, 500 mM NaCl, 1 mM EDTA, and 50 mM Tris pH8. Appropriate caution was exercised when handling acrylamide, a known toxin.

Dissected, fixed, and immunostained samples were treated with AcX stock solution diluted 1:100 in 1xPBS overnight. Brains were then washed with 1xPBS. A gelation chamber was created by applying a Press-to-Seal silicone gasket (ThermoFisher, Cat# P24740) to a glass slide, which was then coated with the PLL solution. AcX-treated brains were immobilized on the PLL surface, up to nine per gasket. Gelation solution was prepared just before gelation, to prevent premature gel polymerization, on ice by adding 10 µL each of 4HT, TEMED, and APS stock solutions to 470 µL of monomer stock solution. Brains were washed with gelation solution and then the gelation chamber was filled with ∼200 µL of gelation solution and incubated on ice for 25 min. The gelation chamber was then sealed with a cover slip and placed in a 37°C incubator to gel and cure for 2 hr.

Gelation chambers were disassembled, and individual gels trimmed close to each brain. Gels were trimmed to a right trapezoid shape to ease specimen orientation. Gels were incubated with proteinase K (NEB, Cat# P8107S) diluted 1:100 (v/v) in proK digestion buffer with shaking, overnight at room temperature. Digested gels were stained in 500 ng/mL DAPI in PBS for 30 min followed by four washes with water for 30min each followed by equilibration overnight. Prepared ExM samples were stored in 1xPBS at 4°C for a week. The samples were expanded in MilliQ water for 3 hours for optimum expansion before imaging by LLSM. Samples expanded ∼4.65x. All samples were scanned within 3-5 hr of expansion.

All ExM samples were imaged in objective scan mode as described (Gao et al., 2019) with minor modifications. For all imaging sessions, focus was maintained by periodic imaging of reference beads. The region of interest was identified by scanning the Brp channel with minimal exposure and acquired as a single tile of 1024×1024×501 voxels.

#### ExLSM sample preparation and imaging

##### Tissue staining and expansion for ExLSM

Tissue for ExLSM were expanded as described above with the following modifications. Fixation and staining of samples for ExLSM was performed with increased concentration of primary and secondary antibodies compared to samples prepared for confocal microscopy (see Table S3). Brains were incubated in gelation solution for 30 minutes at 4°C prior to transfer to the gelation chamber. Digested samples were washed in 1X PBS and stained with 1:1000 DAPI (Sigma Aldrich, Cat# D9542) in 1X PBS for 30 minutes at room temperature, unless antibodies with 405-dye conjugate were used. Samples were expanded to ∼4X in autoclaved milliQ water at room temperature before mounting onto PLL-coated coverslips that were subsequently bonded with Bondic UV-curing adhesive onto a custom fabricated sample holder (Janelia Tech ID 2021- 021) to be suspended vertically in the imaging chamber. Mounted samples were imaged in 1mM Tris Base (Fisher Scientific, Cat# BP152-500) in MilliQ water after a minimum of 2 hours of incubation at room temperature or overnight at 4°C. Unexpanded gels were stored at 4°C in 1X PBS + 0.02% NaN_3_ (Sigma Aldrich, Cat# S8032) for up to 10 days before expansion and imaging.

##### Light-sheet imaging on Zeiss LS7

Images were acquired on a Zeiss LS7 microscope equipped with 405 nm, 488 nm, 561 nm, and 638 nm lasers. Illumination optics with a 10x/0.2 NA were used for excitation (Zeiss, Cat# 400900-9020-000). Detection was performed using a W Plan-Apochromat 20x/1.0 DIC M27 water immersion objective (Zeiss, Cat# 421452-9700-000). The LS7 optical zoom was set to 2.5x, resulting in a total magnification of 50x. CF405S and AF546 dyes were simultaneously excited by the 405 nm and 561 nm laser lines and emission light was separated by a dichroic mirror SBS LP 510 with emission filters BP 420-470 (Zeiss, Cat# 404900-9312-000) and a modified BP 527/23 (Chroma, Cat# ET672/23m). Similarly, AF488 and SeTau647 dyes were simultaneously excited via 488 nm and 638 nm and the emission was split through a dichroic SBS LP 560 with emission filters BP 505-545 and LP 660 (Zeiss, Cat# 404900-9318-000). As the final pair, AF568 and SeTau647 dyes were excited together via the 561 nm and 638 nm laser lines and emission was filtered by a dichroic SBS LP 640 with emission filters BP 575-615 and LP 660 (Zeiss, Cat# 404900-9322-000). To eliminate laser transmission, a 405/488/561/640 laser blocking filter (Zeiss, Cat# 404900-9101-000) was added to the emission path. Images were captured using dual PCO.edge 4.2 detection modules (Zeiss, Cat# 400100-9060-000) with a 50 msec exposure time. Filter and camera alignment were manually calibrated prior to each imaging session. Image volumes were acquired at optimal Z-step and light-sheet thickness and the Pivot Scan feature was used to reduce illumination artifacts by sweeping the light-sheet in the xy-plane. The LS7 microscope was operated using ZEN Black 3.1 (v9.3.6.393).

#### Affinity purification and mass spectrometry

To define GluClα interactors we used affinity purification and mass spectrometry (Figure 5A). To increase confidence of true interactor identification we used two different tagged alleles of GluClα (GluClα-smV5 or GluClα-ALFA) and defined putative interactors as the factors that were identified in mass spectrometry of the immunoprecipitation (IP) with matched nanobody to that of the tag (ALFA or V5) (V5, ProteinTech, Cat# v5tma, ALFA selector, NanoTag, Cat# N1515). As a negative control, we used the same nanobodies against the unmatched tagged allele (e.g., ALFA nanobody with GluClα-smV5 samples). Each set of conditions was done in biological triplicate (Figure 5A). We used stringent criteria to identify GluClα interactors where a putative interactor had to be significantly enriched in both experimental conditions over the negative controls (Figure 5A). In this way, we identified the GluClα interactor Mmd (Table S2).

Immunoprecipitation was performed on 2-10 day old flies of either GluClα-smV5 (w;; GluClα- smV5/+) or GluClα-ALFA (w;; GluClα-1XALFA/+) genotype, which were previously frozen in liquid nitrogen and stored at -80°C. For each pulldown, 10 ml of frozen flies were sieved using a Bel-Art mini-sieve (Bel-Art, Cat# F378451000). Heads were pulverized in liquid nitrogen using a liquid nitrogen chilled mortar and pestle (Fisher Scientific, Cat# FB961C & FB961M) and further homogenized in 1.5 ml of lysis buffer (50 mM Tris HCl (pH 7.6), 150 mM NaCl, 10% glycerol, protease inhibitor (cOmplete mini, Roche, Cat# 11836153001) using a Potter-Elvehjem tissue grinder (Cole Parmer, Cat # EW-04468-14) with 10 strokes at 900 rpm on a Steadystir digital S56 (Fisher Scientific) in a 4°C room on ice, spaced 10s apart to avoid sample overheating. Homogenate was centrifuged at 1,000 xg for 10 min at 4°C to remove the soluble fraction. The insoluble fraction was resuspended in equal volume of lysis buffer with 0.5% sodium deoxycholate (DOC, Sigma-Aldrich, Cat# BCCG2249) and rotated head-over-tail for 30 min at 4°C. Solubilized homogenate was centrifuged for 15 min at 16,000 xg at 4°C. The supernatant was transferred to either V5-Trap magnetic agarose (Chromotek, Cat# V5tma) or ALFA selector PE agarose magnetic beads (NanoTag technologies, Cat# N1515) and washed 2 times in 1 mL of lysis buffer according to the manufacturer protocol. Sample and beads were incubated with head-over-tail rotation for 1 hour at 4°C. Samples were washed with lysis buffer containing 0.5% DOC according to manufacturer’s instructions. Bound protein was eluted in 40 μl of 1X Laemmli buffer (Bio-Rad, Cat# 1610737) for 5 min at 95°C with 1,000 rpm shaking (Fisher Scientific, Cat# 05-400-205).

For proteomic characterization of affinity purified tagged GluClα samples, eluates in 1X Laemmli buffer were diluted two-fold to reduce the SDS concentration. Samples were reduced and alkylated by the sequential addition of 5 mM tris(2-carboxyethyl)phosphine (Goldbio, Cat# 51805-45-9) and 10 mM iodoacetamide (Sigma Aldrich, Cat# I1149). This was followed by protein clean-up using the single-pot, solid-phase-enhanced sample preparation (SP3) protocol (Hughes et al., 2019). Subsequently, the samples underwent proteolytic digestion with Lys-C (NEB, Cat# P8109S) and trypsin (Thermo Scientific, Cat# 90057) at 37°C overnight. The resulting peptides were desalted using SP3-based peptide clean-up and analyzed by LC-MS/MS. Briefly, peptides were separated by reversed phase chromatography using 75 μm inner diameter fritted fused silica capillary column (Polymicro, Cat# 1068150019) packed in-house to a length of 25 cm with bulk 1.9mM ReproSil-Pur beads with 120 Å pores (Dr. Maisch, Cat# r119.aq.) (Jami-Alahmadi et al., 2021). The increasing gradient of acetonitrile (Fisher Chemical, Cat# A955) was delivered by a Dionex Ultimate 3000 (Thermo Fisher Scientific) at a flow rate of 200nL/min. The MS/MS spectra were collected using data dependent acquisition on Orbitrap Fusion Lumos Tribrid mass spectrometer (Thermo Fisher Scientific) with an MS1 resolution (r) of 120,000 followed by sequential MS2 scans at a resolution (r) of 15,000.

#### Co-immunoprecipitation and western blotting

The interaction between GluClα and Mmd was validated by repeating the above immunoprecipitation experiment using magnetic agarose beads with a nanobody recognizing GFP (GFP Selector, NanoTag, Cat# N0315), which interacts with protein harboring the V5 tag (smV5). We used w1118 and w; Rdl-smV5/+ flies as controls to assess specificity of the interaction. IP was carried out as described above on 5 ml of flies and the solubilized homogenate (input) and the eluted protein were run on a denaturing SDS-PAGE using SurePage Bis-Tris 4-20% gels (GenScript, Cat# M00657). Proteins were transferred onto a nitrocellulose membrane with semi-dry transfer (Invitrogen, Cat# PB3210) and membrane were blocked in TBST (10mM Tris-HCl, 0.9% w/v NaCl, 0.02% v/v Tween 20) with 5% skim-milk. Membranes were probed with mouse anti-V5 (SV5-Pk1, 1:500; Abcam Cat# ab27671, RRID:AB_471093), rabbit anti-Mmd, (Guo et al., manuscript under preparation) and mouse anti- αTub (DSHB, Cat# 4A1). Anti-V5 was detected with StarBright Blue 520 goat anti-mouse (Bio-Rad, Cat# 12005867), anti-Mmd with StarBright Blue 700 goat anti-rabbit (Bio-Rad, Cat# 12004162) and anti-αTub with HRP conjugated goat anti-mouse (Jackson ImmunoResearch, Cat#115-036-003). HRP conjugated antibodies were detected with a commercial chemiluminescence system (Millipore, Cat# WBKLS0500) while fluorescent dye conjugated antibodies were directly detected, on an iBright 1500 imaging system (Invitrogen).

#### Secondary antibody labeling

In-house dye conjugation of Fab goat anti-rabbit IgG Fc SeTau647 antibody was performed by cross-linking Fab fragment goat anti-rabbit IgG Fc (Jackson ImmunoResearch, Cat# 111-007- 008, RRID: AB_2632459) with amine-reactive SeTau-647-NHS (SETA BioMedicals, Cat# K9- 4149) at a 1:10 ratio with addition of 0.1M sodium bicarbonate. The mixture was incubated at room temperature with shaking at 500 rpm for 1 hour. Labeled antibody was passed through a Zeba spin desalting columns (ThermoFisher, Cat# 89882). Desalinated labeled antibody was diluted in 40% glycerol at a concentration of 1-5mg/mL and stored at 4°C.

### QUANTIFICATION AND STATISTICAL ANALYSIS

#### EM connectome analysis and visualization

Coordinates of synapses and neuron skeletons for the Fib25 EM dataset were obtained online (https://github.com/janelia-flyem/ConnectomeHackathon2015). Only synapses with a confidence level of 1 were considered in the analysis. For Tm3 neurons, only fully reconstructed neurons (Tm3-home-ant, Tm3-B-ant (rep) and Tm3-C-ant) that fully arborized within the volume in which synapses were annotated were used. For L5 neurons, six L5 neurons (A-F) located in the columns surrounding the central home column, as well as the L5 neuron in the home column, were analyzed. The distribution of synapses from inputs that made three or more synapses onto L5 dendrites in the medulla, was computed in each layer. Layer boundaries were determined using L1 neurons as reference (these neurons arborize in M1 and M5 layers). The neurotransmitter identity of each neuron input was obtained from scSeq data (Kurmangaliyev et al., 2020). The visualization of skeletons and synapses was done using neuTube (Feng et al., 2015).

To visualize T4 and T5 EM data and synaptic inputs, the coordinates of synapses and neuron skeletons for the Fib19 dataset (Shinomiya et al., 2019) were obtained by downloading the data using neuprintr (Bates et al., 2020). This was done for five representative reconstructions of T4 and T5 neurons (Shinomiya et al., 2019). Specifically, for T4 neurons, inputs to the medulla were considered, while for T5 neurons, inputs to the lobula dendrites were analyzed. For Figure 7, all 20 reference T4 dendrites were visualized alongside the Mi1 and Mi9 Home and the ones from columns A-F. Synapses between the home column Mi1 and Mi9 and T4 dendrites were identified. Partners are shown if they made 5 or more synapses, except for a second T4c neuron, which was omitted in Figure 7C for simplicity but formed a similar number of synapses with Mi1 home in its middle domain as seen with T4c-2. The visualization of skeletons and synapses was done using neuTube (Feng et al., 2015).

#### Confocal image analysis

Airyscan processed images of L5, Tm3, T4 and T5 neurons were analyzed using Imaris 9.8. First the membrane channel was extracted using the surface function to create a mask of the neuron. The receptor signal within the mask was analyzed using the spot function to identify puncta above background. All single neuron images shown are 3D projections of masked membrane and receptor signal. To determine the distribution of receptor puncta in T4/T5 dendrites, we restricted the analysis to c/d type T4 or T5 neurons where the direction of dendrite projection was on the long axis of the dendrite, as this facilitated identification of direction of dendritic processes. The proximo-distal axis of dendrites was normalized to 1 and the position of detected puncta was computed onto the normalized axis. All analyses were performed using R Statistical Software (v4.2.2; R Core Team, 2022) using the ggplot2 (Wickham, 2016) package. All other images were processed using FIJI ImageJ (Schindelin et al., 2012).

A Wilcoxon rank sum test with continuity correction was used to obtain significance values when comparing T4, T5, L5 and Tm3 comparison to synapse counts in EM data. Bonferroni correction was applied to p values when comparing L5 and Tm3 synapse counts to EM data in the three distinct dendritic compartments.

#### ExLLSM image analysis

Analysis of ExLLSM images was carried out as previously described (Gao et al., 2019) with minor modifications. All the datasets acquired were deconvolved with the Richardson-Lucy algorithm using experimentally measured point spread functions (PSFs) for each color channel for ten iterations (Chen et al., 2014a). Only the central 360x480x480 voxels where LLSM provides optimal resolution were analyzed. After deconvolution, images were filtered to remove small puncta below 9 voxels by 3D Gaussian kernel with sigma value of 1 and intensity threshold defined as the larger value between the mean plus two standard deviations and the value calculated with Otsu’s method. We applied a lower size threshold (9 instead of 1008 voxels) to include putative non-synaptic receptor signal, that may appear as small puncta, for analysis. The 3D local maxima above the intensity threshold were detected using 3D spheric kernels of 2 voxel-radius by running 3DIJ2 (Haase et al., 2020). The images were segmented using the detected local maxima as seeds to run 3D watershed. The descriptive parameters of segmented objects such as the center of mass (COM) and number of voxels were measured using 3D ImageJ Suite (Ollion et al., 2013). We defined colocalization between receptor and Brp objects if each segmented object of the receptor channel had at least one voxel that was within 1 voxel distance from Brp objects irrespective of distance of the COMs. The Brp-colocalized receptor objects defined by this criterion appeared to be larger and closer to Brp compared to the “non-colocalized” receptor objects (Figures S2M-S2P). The scale shown for ExLLSM and ExLSM images are based on the expansion rate of 4.65, which was estimated by epifluorescence imaging of 16 brains before and after expansion. The expansion rate was larger than previously reported (Gao et al., 2019) presumably due to the use of Glyoxal fixative instead of PFA.

#### ExLSM image analysis

ExLSM images were processed using Python (v3.9.13). Richardson-Lucy deconvolution was performed for 80 iterations on puncta channels stained for nc82 or smV5, and for 10 iterations on membrane channels stained for myr::GFP or myr::tdTomato. Experimental PSFs for deconvolution were acquired by imaging 200 nm fluorescent microspheres (Invitrogen, Cat# T7280) and an average PSF was extracted using the Experimental PSF Wizard in ZEN Blue (v3.4.91.00000). Deconvolved data was intensity normalized in the range of -1 to +40 standard deviations, and binary masks were generated by applying a Laplacian of Gaussian function and setting a positive threshold (0.01 for all myr::GFP and myr::tdTomato channels, 0.02 for all nc82 channels, and 0.1 for all smV5 channels). Puncta channels were then processed as follows. First, local intensity peaks were identified using a 3-voxel diameter window. To group multiple peaks located at the same synapse, nearest neighbors below a defined distance threshold were iteratively averaged, using an 8-voxel radius for Brp-smV5 or nc82 channels, and a 4-voxel radius for GluClα-smV5. The processed peaks were used to perform a marker-controlled watershed transform on the masked image (Meyer and Beucher, 1990), giving the final segmentation results.

Neuron masks were created through manual proofreading and labeling of the membrane channel masks in VVD Viewer (v1.5.10) (for selecting the desired neuron segments) and Napari (v0.4.17) (for manually painting the neuron labels). Nc82 signal within Mi9 axon terminals was extracted by calculating the uniformly weighted centers of mass (i.e., centroids) of each segmented object in the nc82 channel and selecting the objects whose centroids fell within the neuron mask.

All image panels displayed in Figure 6 and Figure S6 show 3D representations produced in VVD Viewer from analyzed data. Each panel shows a representative T4 dendrite from three different brain samples.

#### Analysis of mass spectrometry data

The data generated by LC-MS/MS were analyzed using the MaxQuant bioinformatic pipeline (Cox and Mann, 2008). The Andromeda integrated in MaxQuant was employed as the peptide search engine and the data were searched against Drosophila melanogaster database (Uniprot Reference UP000000803). A maximum of two missed cleavages was allowed. The maximum false discovery rate for peptide and protein was specified as 0.01. Label-free quantification (LFQ) was enabled with LFQ minimum ratio count of 1. The parent and peptide ion search tolerances were set to 20 and 4.5 ppm respectively. The MaxQuant output files were subsequently processed for statistical analysis of differentially enriched proteins using Analytical R tools for mass spectrometry (artMS) (Jimenez-Morales et al., 2021).

#### Sc-RNAseq expression

Heatmaps of expression of genes at 96h APF (Figure S3A) or during pupal development (Figure S5A) was obtained from previously published scSeq RNAseq data (GSE156455, Kurmangaliyev et al., 2020).

## Excel table titles and legends

**Table S3.** Table of genotypes and staining conditions for each figure panel.

