## Supplementary figures and images for "Mapping of multiple neurotransmitter receptor subtypes and distinct protein complexes to the connectome"

### Supplemental Figures

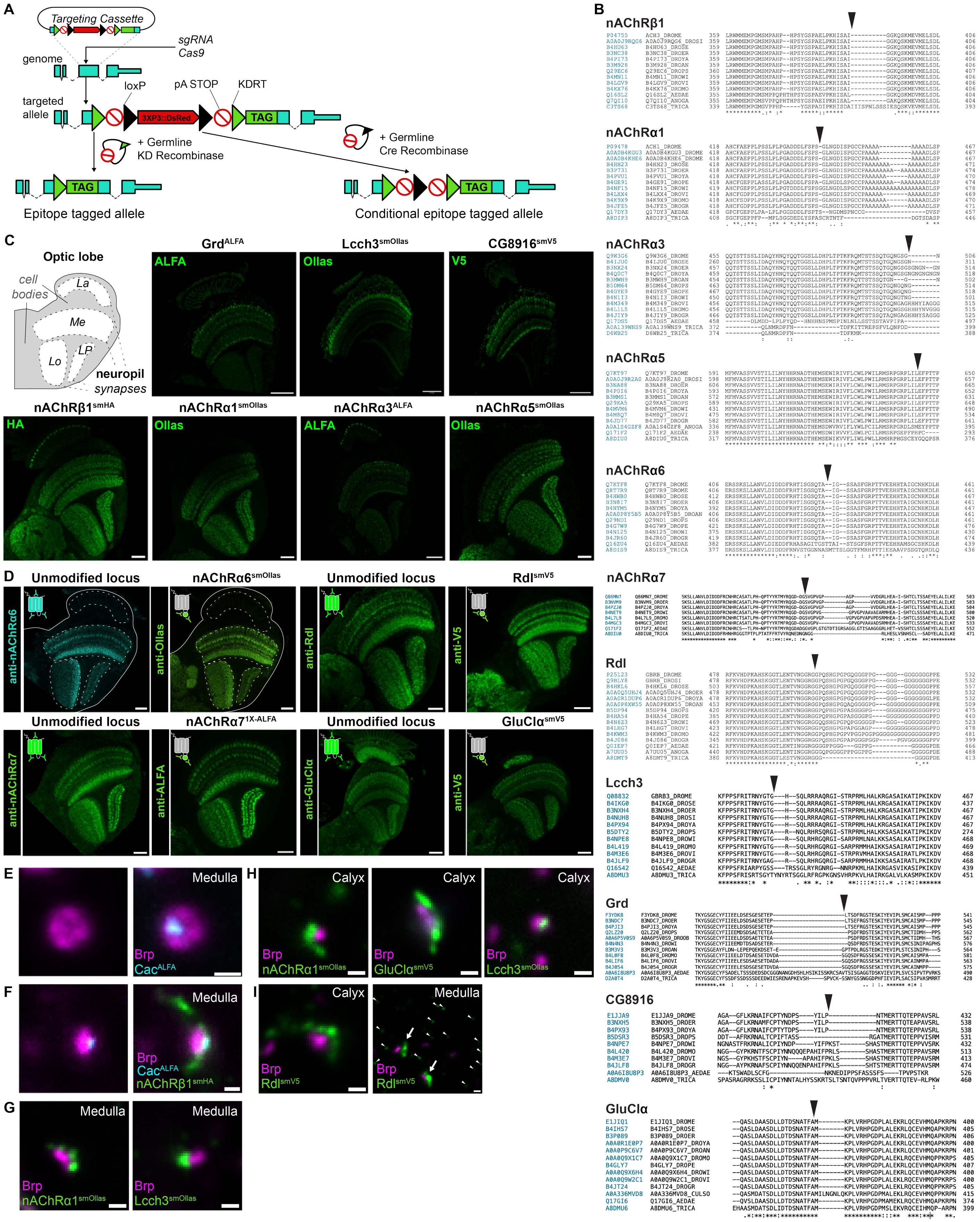

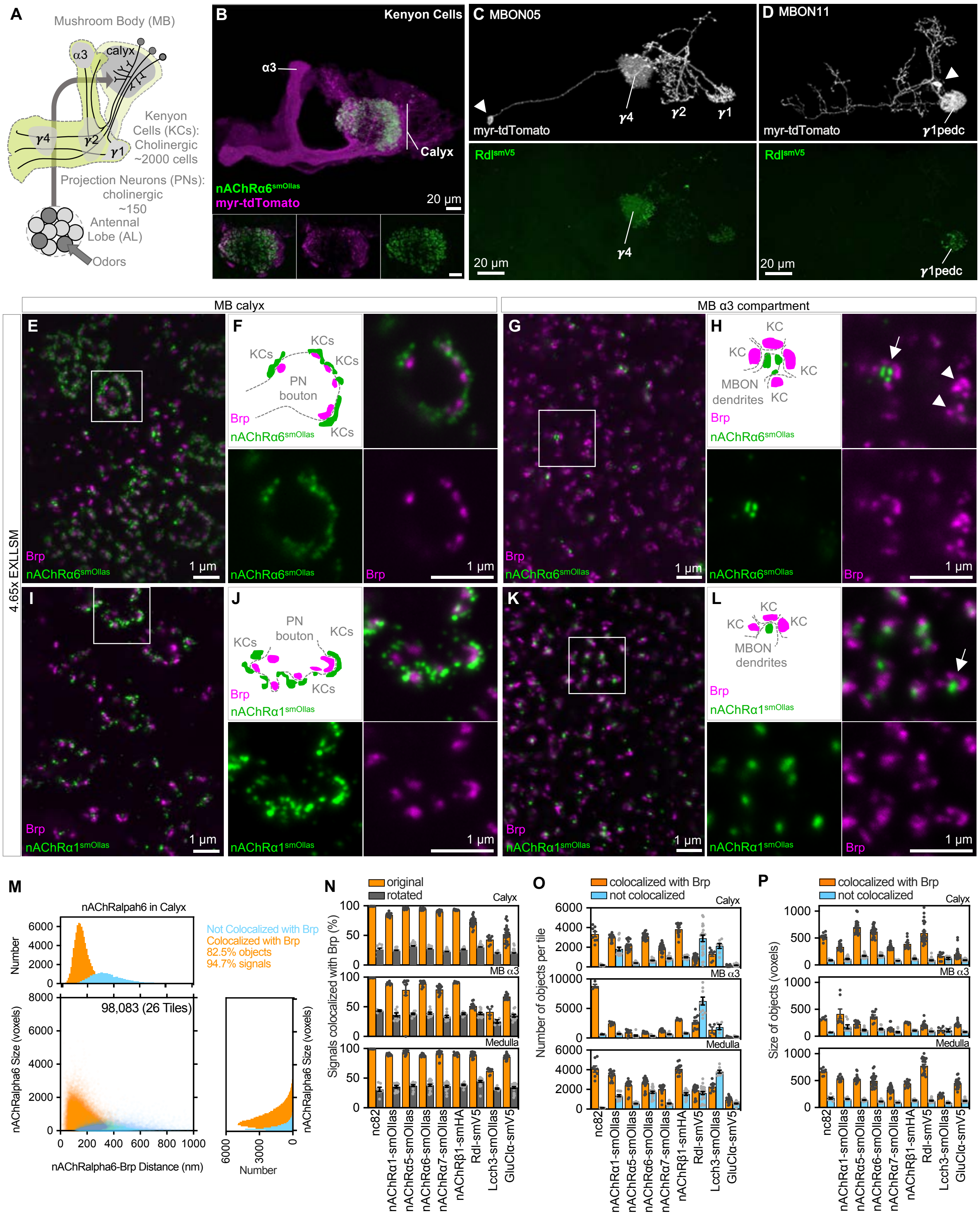

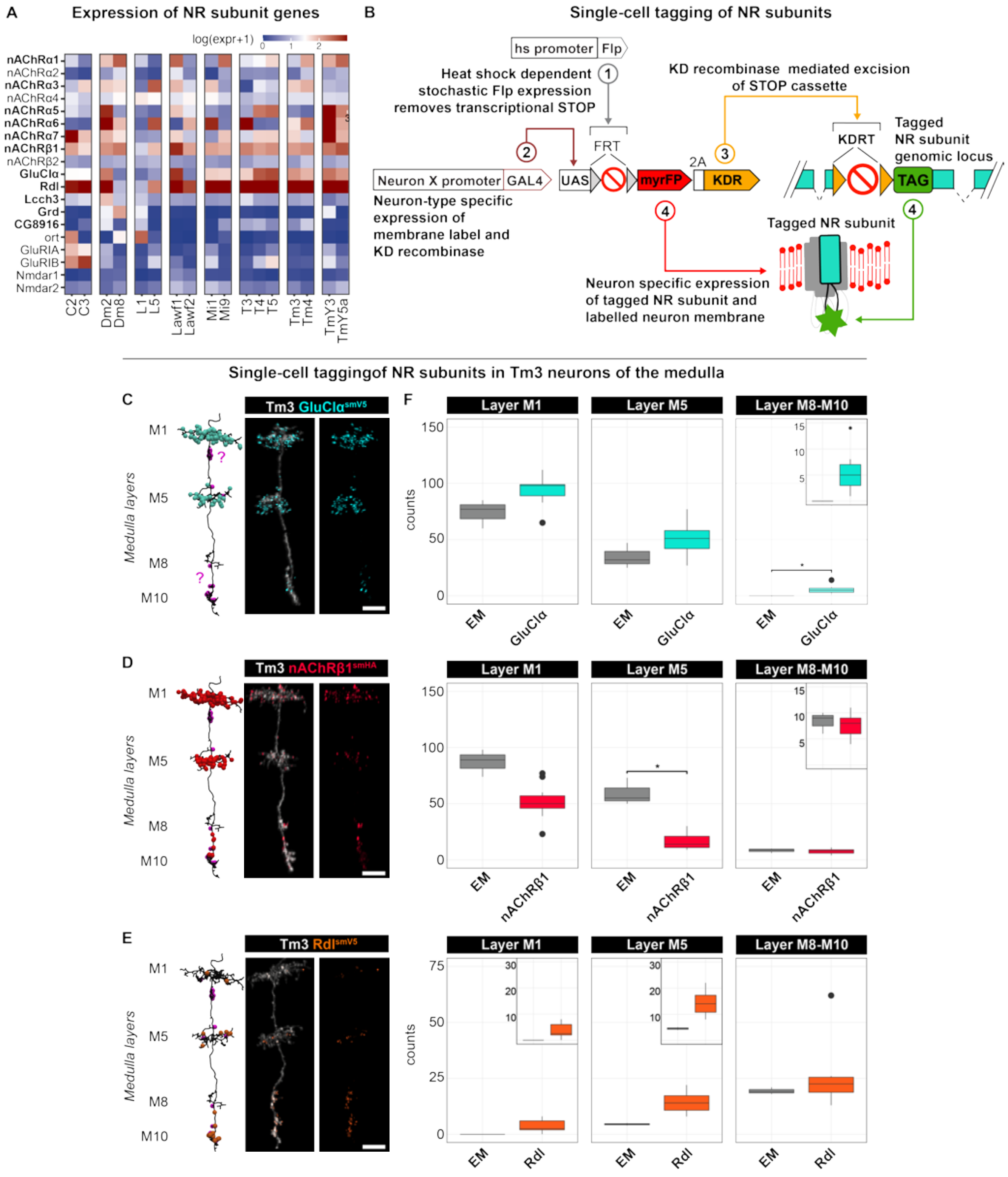

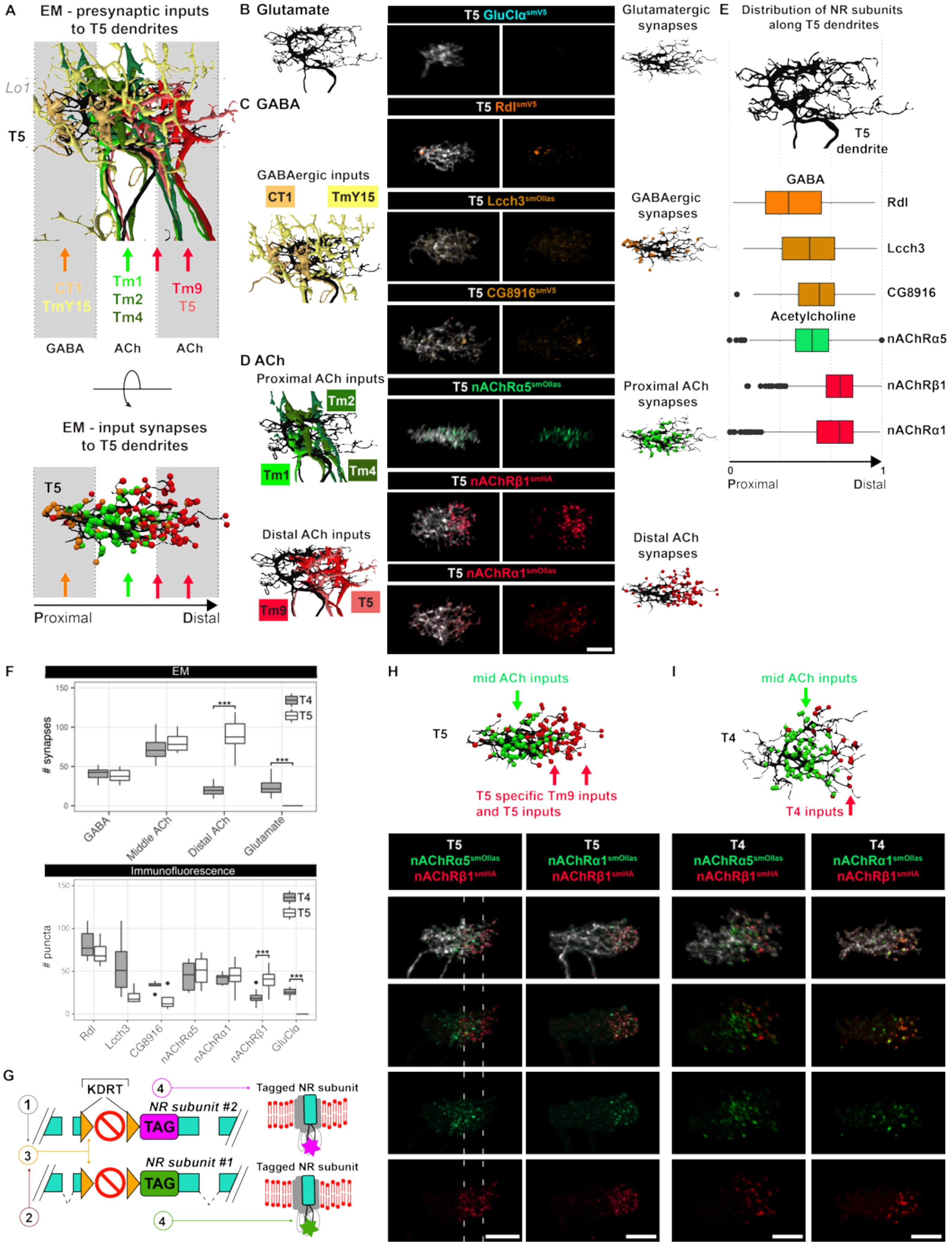

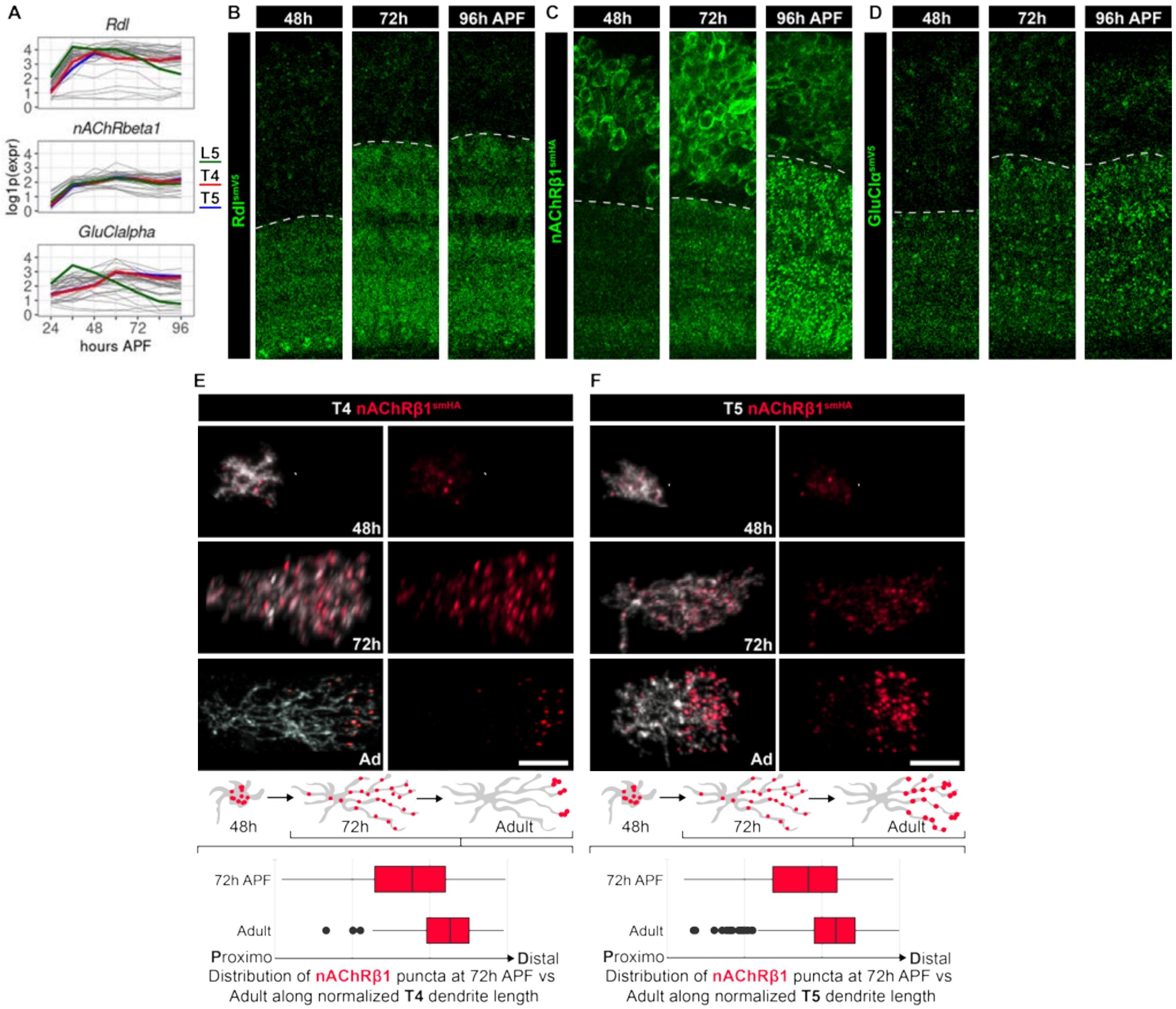

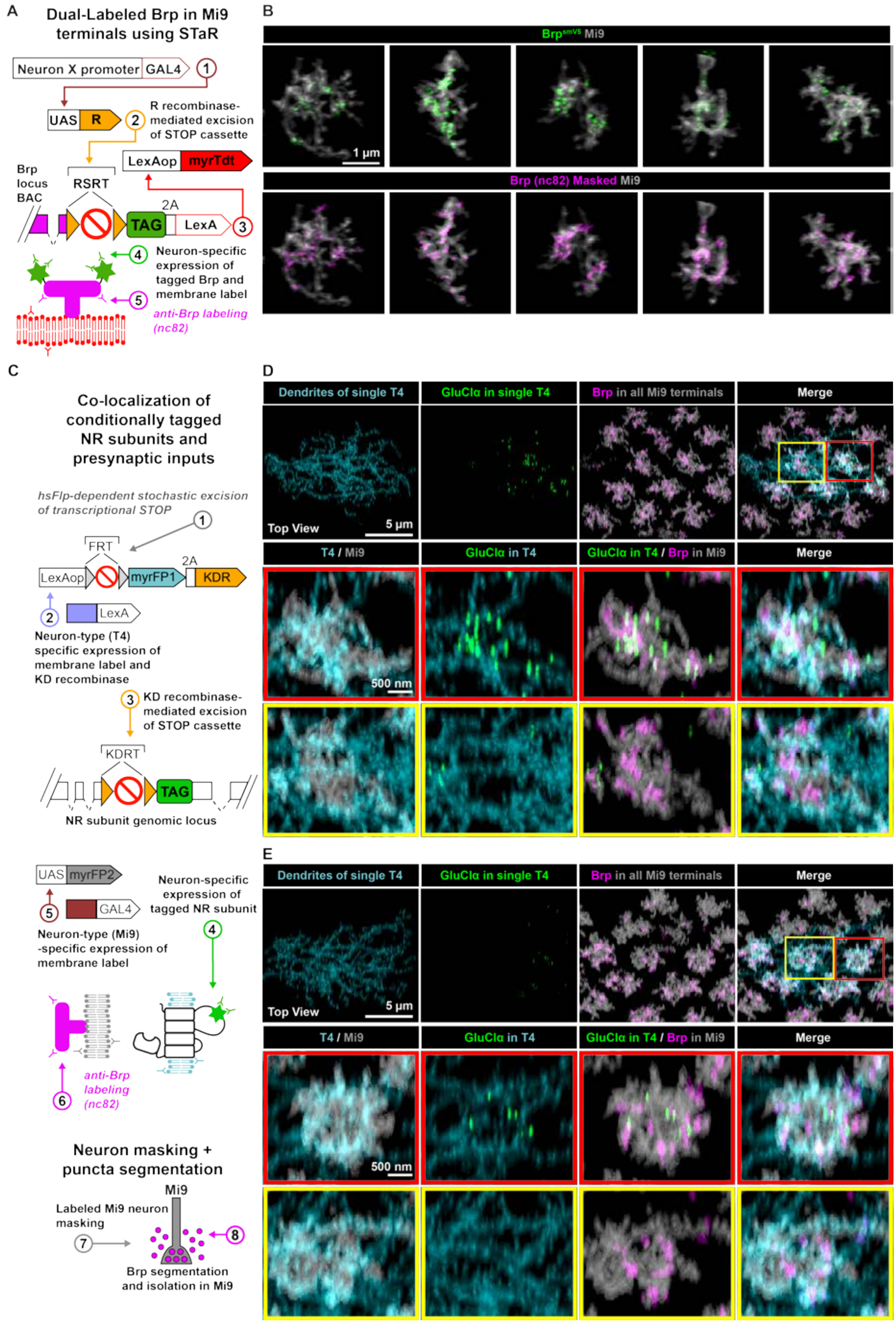
