## Supplementary material for "Mapping of multiple neurotransmitter receptor subtypes and distinct protein complexes to the connectome": Key Resource Table

### Key resources table

| Fly strains (D. melanogaster) | | |
|---|---|---|
| $w^{1118}$ | BDSC | BDSC Cat# 3605; RRID:BDSC_3605 |
| Mmd mutant: Slrp1[1] | BDSC | BDSC Cat# 42249; RRID:BDSC_42249 |
| mmd-V5: w, mmd[Tag:V5.FRT] | BDSC | BDSC Cat# 95303; RRID: BDSC_95303 |
| w;; GluClα-FlpStop.D | Molina-Obando et al., 2019 | |
| w;; nAChRβ1-KDRT-STOP-KDRT-smGdP-10xHA/TM6B | This study | |
| w;; nAChRα1-KDRT-STOP-KDRT-smGdP-10xOllas/TM6B | This study | |
| w, nAChRα3-KDRT-STOP-KDRT-1xALFA;; | This study | |
| w; nAChRalpha5-KDRT-STOP-KDRT-smGdP-10xOllas/CyO; | This study | |
| w; nAChRalpha6-KDRT-STOP-KDRT-smGdP-10xOllas/CyO; | This study | |
| w, nAChRα7-KDRT-STOP-KDRT-1xALFA;; | This study | |
| w, nAChRα7-KDRT-STOP-KDRT-smGdP-10xOllas;; | This study | |
| w;; Rdl-KDRT-STOP-KDRT-smGdP-10xV5/TM6B | This study | |
| w;; Rdl-KDRT-STOP-KDRT-smGdP-10xHA/TM6B | This study | |
| w;; Rdl-KDRT-STOP-KDRT-1xALFA/TM6B | This study | |
| w, Lcch3-KDRT-STOP-KDRT-smGdP-10xOllas;; | This study | |
| w;; Grd-KDRT-STOP-KDRT-1xALFA | This study | |
| w, CG8916-KDRT-STOP-KDRT-smGdP-10xOllas;; | This study | |
| w;; GluClα-KDRT-STOP-KDRT-smGdP-10xV5/TM6B | This study | |
| w;; GluClα -KDRT-STOP-KDRT-1xALFA/TM6B | This study | |
| w;; nAChRβ1-KDRT-smGdP-10xHA | This study | |
| w;; nAChRα1-KDRT-smGdP-10xOllas | This study | |
| w, nAChRα3-KDRT-1xALFA;; | This study | |
| w; nAChRalpha5-KDRT-smGdP-10xOllas; | This study | |
| w; nAChRalpha6-KDRT-smGdP-10xOllas; | This study | |
| w, nAChRα7-KDRT-1xALFA;; | This study | |
| w, nAChRα7-KDRT-smGdP-10xOllas;; | This study | |
| w;; Rdl-KDRT-smGdP-10xV5/TM6B | This study | |
| w;; Rdl-KDRT-smGdP-10xHA/TM6B | This study | |
| w;; Rdl-KDRT-1xALFA/TM6B | This study | |
| w, Lcch3-KDRT-smGdP-10xOllas;; | This study | |
| w;; Grd-KDRT-1xALFA | This study | |
| w, CG8916-KDRT-smGdP-10xOllas;; | This study | |

| | | |
|---|---|---|
| w;; GluClα-KDRT-smGdP-10xV5/TM6B | This study | |
| w;; GluClα -KDRT-1xALFA/TM6B | This study | |
| w;; P{w+, 13xLexAoP-FRT-STOP-FRT-myrGFP-2A-KDR.PEST}attP1 | This study | |
| w, P{w+, 10xUAS-FRT-STOP-FRT-myrGFP-2A-KDR.PEST}attP40 | This study | |
| P{w+, 10xUAS-FRT-STOP-FRT-myrTdt-2A-KDR.PEST}attP5 | This study | |
| w, 10xUAS-myrTdtomato;; | BDSC | BDSC Cat# 32223;<br>RRID: BDSC_32223 |
| w, Cac-1xALFA;; | Bhukel et al., manuscript in preparation | |
| w, P{w+, hsFLPG5.PEST}attP3;; | BDSC | BDSC Cat# 62118;<br>RRID: BDSC_62118 |
| w, P{hs-FLPG5.PEST.Opt)attP3;; | BDSC | BDSC Cat# 77140;<br>RRID: BDSC_77140 |
| PBac{brp-FRT-STOP-FRT-smGdP-10xV5-2A-LexA-VP16}VK00018 | laboratory stock | |
| MBON-γ1pedc>α/β: sGal4$^{MB112C}$:<br>w;; P{R13F04-GAL4.DBD}attP2,<br>PBac{R93D10-p65.AD}VK00027 | BDSC | BDSC Cat# 68325;<br>RRID: BDSC_68325 |
| MBON-γ4>γ1γ2, MBON-β1>α: sGal4$^{MB434B}$:<br>w; P{R30E08-p65.AD}attP40/CyO;<br>P{R53C10-GAL4.DBD}attP2 | BDSC | BDSC Cat# 68325;<br>RRID: BDSC_68325 |
| Kenyon Cells sGal4$^{MB010C}$:<br>w; P{R13F02-p65.AD}attP40;<br>P{R52H09-GAL4.DBD}attP2 | BDSC | BDSC Cat# 68293;<br>RRID: BDSC_68293 |
| MBONα3, MBONα′2 sGal4$^{MB082C}$:<br>w;; P{R23C06-GAL4.DBD}attP2,<br>PBac{R40B08-p65.AD}VK00027 | BDSC | BDSC Cat# 68286;<br>RRID: BDSC_68286 |
| L5-gal4$^{R64B07}$: w;; P{w+, GMR64B07-Gal4}attP2 | BDSC | BDSC Cat# 39293;<br>RRID: BDSC_39293 |
| Tm3-gal4$^{R13E12}$: w;; P{GMR13E12-Gal4}attP2 | BDSC | BDSC Cat# 48569;<br>RRID: BDSC_48569 |
| T4T5-gal4$^{R42F06}$: w;; P{GMR42F06-Gal4}attP2 | BDSC | BDSC Cat# 41253;<br>RRID: BDSC_41253 |
| T4T5-LexA$^{R42F06}$: w; P{w+, GMR42F06-LexA}attP40 | BDSC | BDSC Cat# 54203;<br>RRID: BDSC_54203 |
| Mi9 sGal4$^{SS02432}$:<br>w; P{R48A07-p65.AD}attP40;<br>P{VT046779-GAL4.DBD}attP2 | BDSC | BDSC Cat# 86854;<br>RRID: BDSC_86854 |
