## Supplementary material for "Mapping of multiple neurotransmitter receptor subtypes and distinct protein complexes to the connectome": Table S1

| **Gene** | **Reference protein isoform^a^** | **Insertion site^b^** | **Insertion location^c^** | **Tags^d^** | **gRNA-target(PAM)** |
| --- | --- | --- | --- | --- | --- |
| nAChRβ1 | FBpp0073155 | I391 | chr3L:4434861 | smHA | CTTGGATTGCTTGCCGCCGA(TGG) |
| nAChRα1 | FBpp0084003 | S443 | chr3R:24403383 | smOllas | ATGTCTCCGTTGAGGCCCGA(CGG) |
| nAChRα3 | FBpp0288426 | G505 | chrX:8285689 | 1XALFA | AGACGGGTCAGAATGGCAGC(GGG) |
| nAChRα5 | FBpp0089367 | L644 | chr2L:14088048 | smOllas | AGGGCGTGGTCGGGAACTCC(AGG) |
| nAChRα6 | FBpp0079505 | A404 | chr2L:9797925 | smOllas | TTACGCCGACGAGCCAATGG(CGG) |
| nAChRα7 | FBpp0089248 | G464 | chrX:19330238 | smOllas  1XALFA | GTGACGGCAGCGTGGGACCCG(TGG) |
| Rdl | FBpp0076261 | R501 | ch3L: 9151410 | smV5  smHA  1XALFA | ACGGTGAATGGCGGACGCGG(TGG) |
| Lcch3 | FBpp0089263 | T427 | chrX: 15928138 | smOllas | ACGTCTAAGCTGGCTATGTC(CGG) |
| Grd | FBpp0074905 | P504 | chr3L:17838128 | 1XALFA | AGCCACTGACCTCGGACTTT(CGG) |
| CG8916 | FBpp0290450 | P517 | chrX:15924237 | smV5 | CGTTTGGGTGGTTCTCTCCA(TGG) |
| GluClα | FBpp0307401 | A375 | chr3R:19765748 | smV5 | TAGCAATGCAACGTTCGCAA(TGG) |
| ^a^ Flybase protein isoform reference for aminoacid insertion site.  ^b^ Targeted exon is common to all annotated protein isoforms (FlyBase r. FB2023_01).  ^c^ Coordinates based on dm6 *D. melanogaster* reference genome.  ^d^ smX – smGdP – spaghetti monster Green darkened Protein tagged with 10 copies of the indicate epitope tag. | | | | | |

**List of tagged NR subunits and insertion sites**
