## Supplementary material for "Mapping of multiple neurotransmitter receptor subtypes and distinct protein complexes to the connectome": Table S2

| **Protein** | **GeneName** | **Comparison** | **log2FC** | **pvalue** | **adj.pvalue** | **imputed** | **iLog2FC** | **iPvalue** |
| --- | --- | --- | --- | --- | --- | --- | --- | --- |
| E1JIQ1 | GluClalpha | nbV5::GluClalpha-smV5 vs nbV5::GluClalpha-1XALFA | #NUM! | – | 0 | yes | 6.31E+00 | 3.79E-02 |
| Q9VXL1 | mmd | nbV5::GluClalpha-smV5 vs nbV5::GluClalpha-1XALFA | #NUM! | – | 0 | yes | 6.07E+00 | 3.52E-02 |
| E1JIQ1 | GluClalpha | nbALFA::GluClalpha-1XALFA vs nbALFA::GluClalpha-smV5 | 7.89E+00 | 5.30E-05 | 1.17E-02 | no | 7.89E+00 | 1.17E-02 |
| M9NH46 | mmd | nbALFA::GluClalpha-1XALFA vs nbALFA::GluClalpha-smV5 | #NUM! | – | 0 | yes | 3.15E+00 | 1.78E-02 |
| Q9VXL1 | mmd | nbALFA::GluClalpha-1XALFA vs nbALFA::GluClalpha-smV5 | 4.90E+00 | 4.24E-06 | 1.88E-03 | no | 4.90E+00 | 1.88E-03 |

### Putative GluClα interactors as identified by AP-MS
